# An integrated model of population growth saturation and basal ROS levels predicts cellular ferroptosis sensitivity

**DOI:** 10.1101/2025.09.15.676259

**Authors:** Eric Seidel, E. Yaren Itak, Fabienne Müller, F. Isil Yapici, Johannes Brägelmann, Johannes Berg, Silvia von Karstedt

## Abstract

Ferroptosis is a form of regulated cell death characterized by iron-dependent lipid peroxidation and membrane rupture. While cellular populations reaching confluence are known to have limited sensitivity to ferroptosis, an understanding of the interplay between growth dynamics, ROS levels and ferroptosis is currently lacking. Here we use live-cell imaging coupled to ROS tracing to reveal a feedback loop between population growth and ferroptotic cell death. Starting out from the observation that the cellular proliferation rate declines with increased cellular density, we find that ROS levels also decline with increasing cellular density. Low ROS levels make cells insensitive to ferroptosis, which in turn enables population growth. This feedback produces two steady states: (i) a ferroptosis-insensitive state characterized by slow growth, low levels of ROS and low rates of cell death and (ii) a ferroptosis-sensitive state characterized by rapid growth, ROS accumulation, and high rates of ferroptosis. Interestingly, triggering effective ferroptosis by interfering with GPX4 activity is directly linked with this mechanism. On the other hand, keeping cell numbers and drug concentration/cell constant while restricting growth space led to reduced proliferation, reduced ROS and decreased ferroptotic cell death. Importantly, ferroptosis resistance at high cellular confluency could be broken by increasing cellular ROS and lipid ROS through a galactose-promoted OXPHOS switch. A mathematical model of the feedback mechanism predicts the long-term fate of populations as well as their ferroptosis sensitivity when external conditions impacting cell proliferation rates, ROS, or both are changed.

## Introduction

The observation that cysteine removal kills cells in culture only under low density dates back to the establishment of human cell culture by Harry Eagle in the 1960ies(Eagle, 1960). We now know that in cell culture cysteine removal induces ferroptosis, an iron-dependent type of regulated necrosis(Gao et al., 2018). Ferroptotic cells present with loss of plasma membrane integrity due to excessive lipid peroxidation, osmotic cytoplasmic swelling, and mitochondrial fragmentation and failure(Dixon et al., 2012; Friedmann Angeli et al., 2014; Riegman et al., 2020). Glutathione peroxidase 4 (GPX4) constitutively hydrolyses lipid hydroperoxides thereby preventing ferroptosis(Yang et al., 2014). Consequently, genetic or pharmacological inactivation of GPX4 induces ferroptotic cell death(Friedmann Angeli et al., 2014; Seiler et al., 2008; Yang et al., 2014). GPX4 requires glutathione (GSH) as an electron donor to reduce lipid hydroperoxides. Cellular GSH synthesis is coupled to the availability of intracellular cysteine which can be generated from cystine imported from the extracellular space via the cystine/glutamate antiporter System xc-. One of the most defining hallmarks of ferroptotic cell death to date is excessive membrane lipid peroxidation driven by lipid reactive oxygen species (lipid ROS) which unlike general ROS reside within cellular membranes(Kagan et al., 2017; Wiernicki et al., 2020). Yet, in cycling cells mitochondria are a major source of superoxide as a by-product of oxidative phosphorylation (OXPHOS)(Ivanova et al., 2021), which in the presence of protein iron-sulphur clusters undergoes a Fenton reaction giving rise to lipid ROS(Homma et al., 2021). Thereby, mitochondrial OXPHOS is thought to indirectly feed into ferroptosis susceptibility of cells. Notably, cells generating cellular ATP mainly through glycolysis—the so-called Warburg effect—also generate less superoxide (Martens et al., 2005), which may influence ferroptosis sensitivity(Homma et al., 2021). Unlike for other types of cell death, cellular population density indeed has been observed to also restrict the capability of cells to undergo ferroptotic cell death induced through various experimental means(Schneider et al., 2010; Seiler et al., 2008). Herein, E-cadherin on epithelial cells was shown to sequester YAP1 upon close cell-to-cell contact thereby preventing YAP1-mediated induction of ACSL4 and TFRC(Wu et al., 2019). Yet, similar observations can also be made for mesenchymal cells lacking expression of E-cadherin suggesting additional mechanisms by which high cellular confluence restricts ferroptotic cell death (reviewed in(Vucetic et al., 2020)). Indeed, high cellular density was shown to downregulate ROS and ferric iron content(Yan et al., 2024), known outputs and inputs, respectively, of mitochondrial oxidative phosphorylation (OXPHOS). While the mitochondrial TCA cycle was shown to promote ferroptosis induced through cysteine or GSH depletion(Gao et al., 2018), it is only poorly understood how cellular confluence, cellular ROS levels and ferroptosis sensitivity might be connected. Here, we used a combined approach of cellular experimental work with mathematical modelling to address this connection.

## Results

### Population growth dynamics are linked with cellular ROS levels

To test the hypothesis that cellular proliferation rates are accompanied by high metabolic generation of ROS in non-epithelial cells, we made use of mouse embryonic fibroblasts (MEFs) seeded at low (LD), middle (MD) and high cellular density (HD) into similar sized cell culture wells in the same volume of growth media, tracked cellular confluence over time using live cell imaging and calculated doubling times. As expected, cells seeded at HD reached confluence first, followed by MD and LD (Figure 1A). Importantly, cells seeded at HD showed progressively slower proliferation rates in comparison to cells seeded at LD or MD (Figure 1B). Next, we measured levels of cellular ROS comparing LD, MD and HD cells using the fluorescent ROS tracer dye CellROX. Indeed, mean fluorescent intensity (MFI) of the tracer decreased in all three starting densities over time with increased cellular density indicative of lower levels of ROS (Figure 1C). Notably, with labile iron present, lipid ROS can arise through Fenton reactions. Indeed, basal cellular lipid ROS levels also decreased inversely proportional with starting density (Figure 1D). To next determine whether these changes in cellular proliferation and ROS levels were mirrored in transcriptional profile changes, we subjected MEFs seeded at LD as compared to HD into similar sized wells with similar volumes of media to RNA-sequencing (RNA-seq). Interestingly, mere changes in growth at these two different seeding densities was sufficient to significantly alter gene expression programmes in the two states. Interestingly, gene set enrichment analysis (GSEA) showed an enrichment of mitotic spindle gene set as well as ROS pathway (Figure 1E), suggesting proliferative cells with high levels of ROS to be present predominantly within the LD condition.

**Figure 1.**
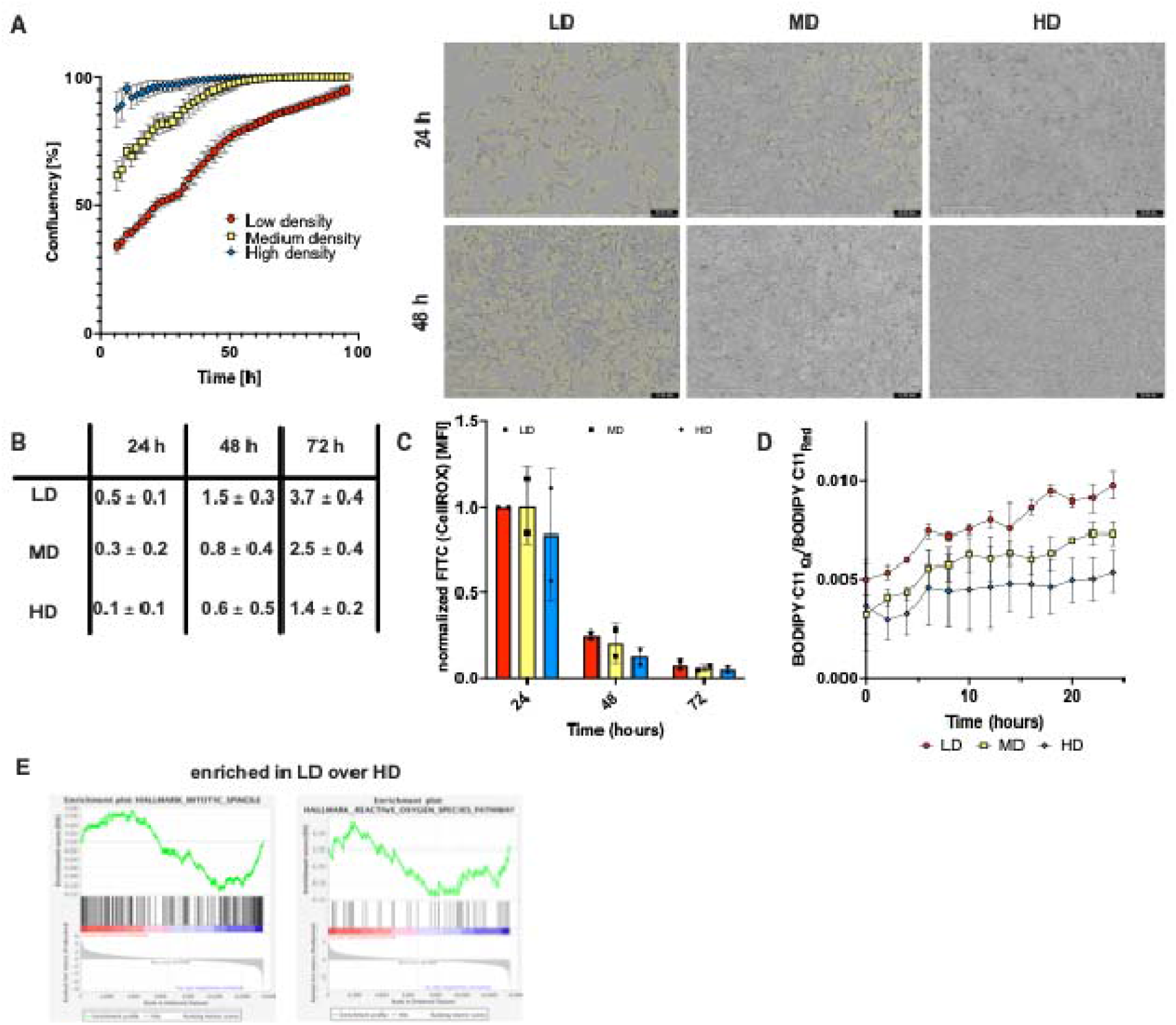
Population growth is linked to cellular ROS levels. (A) Confluency over time measured in MEFs seeded at 25.000 (low density, LD), 55.000 (medium density, MD) or 120.000 cells (high density, HD) per well on a 24 well plate using the Incucyte live cell imaging system. Right: brightfield images recorded at the indicated time points. Yellow margins indicate cell boundaries. (B) Population doublings of MEF populations seeded at LD, MD and HD measured at the indicated time points. (C) Total cellular ROS measured using the CellROX dye in FACS at the indicated time points of MEF populations seeded at LD, MD and HD. Shown is MFI(mean fluorescence intensity) normalized to LD at 24 h within each of the replicates. (D) Lipid ROS over time shown as oxidized-to-reduced ratio of the STY-BODIPY dye, measured in the Incucyte live-cell imaging system at the indicated time points of MEF populations seeded at LD, MD and HD. (E) Exemplary gene sets from Gene set enrichment analysis (GSEA) are shown.

### A mathematical model predicts a negative feedback loop of population growth, lipid ROS and ferroptosis

Given that high lipid ROS at LD is likely permissive for ferroptotic cell death, we decided to mathematically model a connection of these parameters to understand mechanisms potentially interlinking these. To this end, we built a minimal mathematical model of the interaction between population growth, levels of lipid ROS, and ferroptosis. The model consists of three components: i) population dynamics driven by the balance between logistic growth and ferroptotic cell death, ii) the production of ROS as a side-product of cellular growth, which can feed into to the production of lipid ROS and, via the Fenton reaction, to lipid peroxidation (LPO) and iii) ferroptotic cell death triggered by LPO. Interestingly, the model predicts a negative feedback loop in which fast growth leads to high levels of lipid ROS, which via LPO production can lead to ferroptotic cell death in turn restricting growth (Figure 2A). We focus on the population size n and average levels of lipid ROS per cell, denoted r. The dynamics of these variables is characterized by two differential equations: The population size grows logistically, reaching a plateau specified by a carrying capacity K. It decreases due to cell death, which effectively occurs when lipid ROS reaches a level r0 such that lipid ROS-dependent LPO production overwhelms the LPO scavenging and repair mechanisms and cells undergo ferroptosis. Lipid ROS levels increase at a rate a per cell division and they decrease at a rate br due to ROS scavenging and degradation (Figure 2B). We also included a recently described biochemical positive-feedback loop, where the generation of lipid ROS leads to the production of further lipid ROS as a by-product of the Fenton reaction(Co et al., 2024). The resulting differential equations (Figure 2B) predict the dynamics and the eventual fate of a population (Figure 2C). Interestingly, the feedback between population dynamics, lipid ROS levels and ferroptotic cell death leads to seeding-density-dependent cell fates. Each point in this diagram specifies a lipid ROS level r on the x-axis and a population size n on the y-axis. Arrows indicate the flow of populations over time: Starting from low population sizes (relative to the carrying capacity) and high levels of lipid ROS, populations are taken to a steady state with high rate of cell death and low population size. This ferroptosis-sensitive state is indicated by a red point. Conversely, initial conditions with high population size and low levels of lipid ROS evolve to a steady state characterized by a low rate of ferroptotic cell death and a population reaching carrying capacity. Initial conditions that lead to either the susceptible or the resistant state (i.e. HD, MD or LD seeding of cells or varying levels of cellular growth and, hence ROS production) define basins of attraction of these states (shaded in blue and red, respectively). These basins of attraction depend on the parameters of the model (see Document S1).

**Figure 2.**
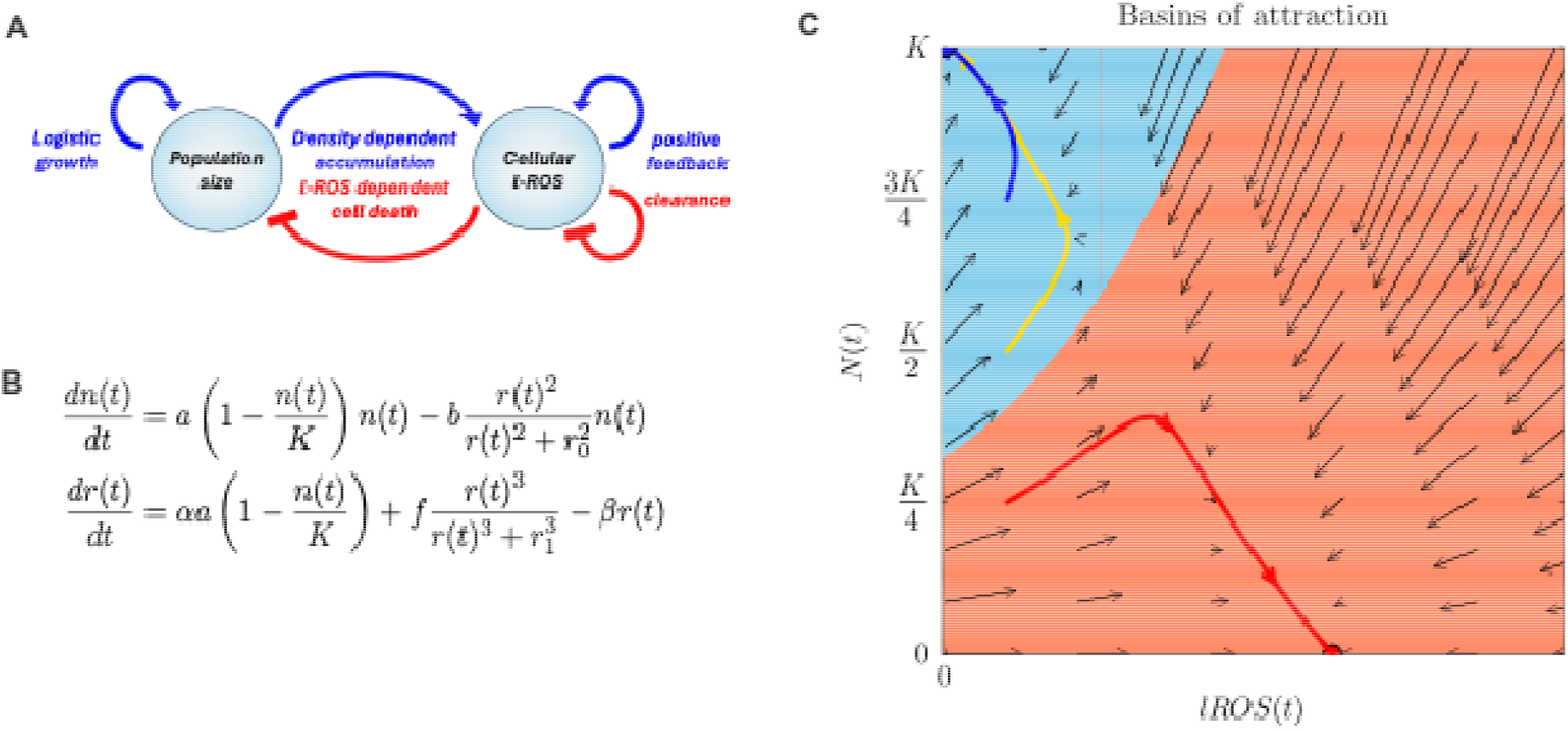
A mathematical model of the interaction between population growth, levels of lipid ROS, and ferroptosis. (A) Schematic overview of the feedback loop linking population dynamics and lipid ROS levels. (B) Mathematical model describing the feedback loop linking population dynamics and lipid ROS levels. n, population size; r, average levels of lipid ROS per cell; K, carrying capacity; a basal rate of cell divisions; b basal rate of cell death; a, ROS generation rate per cell division; b, ROS depletion rate per molecule. (C) Basin of attraction plot resulting from the mathematical model, where the lines represent sample trajectories starting from different seeding densities (blue – HD; yellow – MD; red – LD). Areas colored orange or blue indicate basins on attraction, i.e. starting points that will evolve either to the ferroptosis sensitive state (orange) or the insensitive state (blue). Arrows indicate the dynamics defined by the model (see Document S1).

### Seeding density determines cellular ROS levels and ferroptosis sensitivity

To test our mathematical model experimentally, we manipulated the parameters characterizing the feedback loop between population dynamics, ROS levels and ferroptosis. We started by interfering with GPX4 activity using the small molecule inhibitor RSL3(Yang and Stockwell, 2008). Since constitutive GPX4 activity leads to clearance of LPO products, GPX4 inhibition leads to increased levels of LPO. Excessive LPO leads to membrane destabilization, permeabilization(Pedrera et al., 2020) and ferroptotic cell death(Bebber et al., 2020; Conrad and Pratt, 2019). Interestingly, RSL3 treatment led to increased detection of both lipid ROS and total ROS as the seeding cell density was decreased from HD to LD (Figure 3A, 3B). Moreover, lipid ROS increased over time in a strict density-dependent manner upon RSL3-treatment (Figure 3C). As a result of this, cells seeded at LD and MD but not HD underwent cell death upon RSL3 treatment (Figure 3D). Importantly, these outcomes match the predictions of our mathematical model: We fit the model parameters to the experimental data giving confluence and lipid ROS levels over time, resulting in the curves shown (Figures 3E-3J), both for the control without RSL3 and the experiments with GPX4 inhibition using RSL3. As expected, we find the lipid ROS threshold parameter r0 reduced under RSL3 relative to the control, which leads to a shift in the basins of attraction specifying cell fate (Figure 3G, 3J): under RSL3, cell death is triggered already at lower lipid ROS rates due to reduced LPO clearance. This reduces the population size, and leads to a faster turnover of cells, and (averaged over the surviving cells) increased levels of lipid ROS.

**Figure 3.**
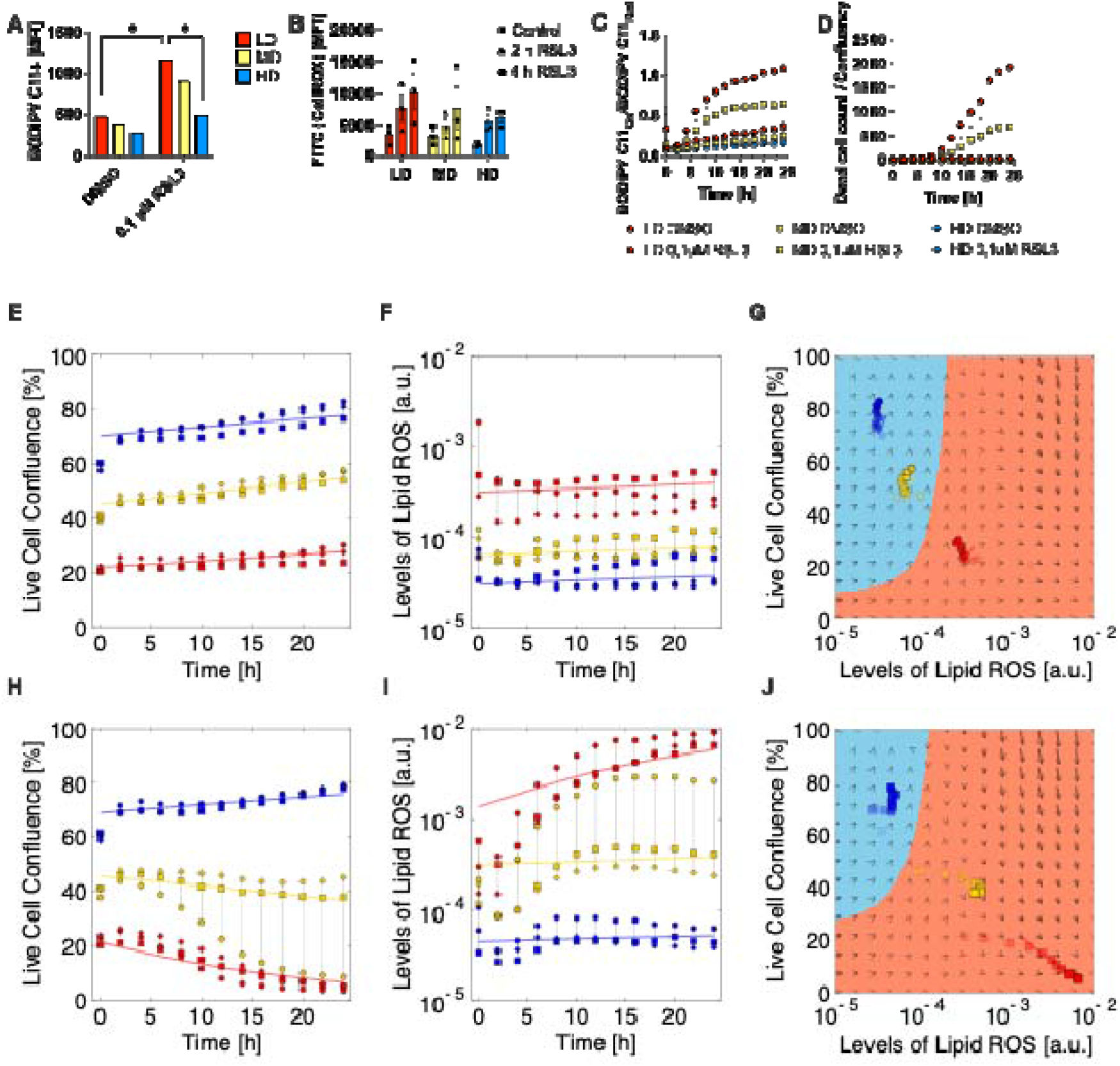
Ferroptosis sensitivity is linked with the clearance of lipid peroxidation products. (A) Total lipid ROS measured using Bodipy C11 dye in FACS at the indicated time points of MEF populations seeded at LD, MD and HD in 24-well plates and in presence of vehicle control (DMSO) or GPX4 inhibitor 0.1 µM RSL3. MFI, mean fluorescence intensity. (B) Total cellular ROS measured using the CellROX dye in FACS at the indicated time points of MEF populations seeded at LD, MD and HD in 12-well plates and in presence of DMSO or 0.1 µM RSL3. (C) Lipid ROS over time shown as oxidized-to-reduced ratio of the STY-BODIPY dye, measured in the Incucyte live-cell imaging system at the indicated time points of MEF populations seeded at LD, MD and HD in 12-well plates and in presence of DMSO or 0.1 µM RSL3. (D) Dead cell counts over time shown as dead cell count-to-confluence ratio using t DRAQ7 dead cell dye, measured in the Incucyte live-cell imaging system at the indicated time points of MEF populations seeded at LD, MD and HD in 12-well plates and in presence of DMSO or 0.1 µM RSL3. Data in A-D are shown as mean ± SEM.; * p ≤ 0.05 (one-way Anova). (E) Live cell confluency of the Incucyte measurement corresponding to data shown in (C)+(D), DMSO-treated populations. Colors indicate different starting population sizes (blue – HD; yellow – MD; red – LD), and differently shaped symbols represent different replicates. Data from DMSO-treated populations shown. (F) Levels of lipid ROS in live cells computed from the Incucyte measurements displayed in (C)+(D), DMSO-treated population. Colors indicate different starting population sizes (blue – HD; yellow – MD; red – LD), and differently shaped symbols represent different replicates. (G) Basins of attractions plot showing live cell confluence versus live cell lipid ROS levels computed from (E) and (F). Colors of symbols indicate different starting population sizes (blue – HD; yellow – MD; red – LD), for clarity only a single replicate is shown. Data from DMSO-treated populations. (H) Live cell confluency of the Incucyte measurement corresponding to data shown in (C)+(D), DMSO-treated populations. Colors indicate different starting population sizes (blue – HD; yellow – MD; red – LD), and differently shaped symbols represent different replicates. Data from 0.1 µM RSL3-treated populations shown. (I) Levels of lipid ROS in live cells computed from the Incucyte measurements displayed in (C)+(D), DMSO-treated population. Colors indicate different starting population sizes (blue – HD; yellow – MD; red – LD), and differently shaped symbols represent different replicates. Data from 0.1 µM RSL3-treated populations. (J) Basins of attractions plot showing live cell confluence versus live cell lipid ROS levels computed from (H) and (I). Colors of symbols indicate different starting population sizes (blue – HD; yellow – MD; red – LD), for clarity only a single replicate is shown. Data from 0.1 µM RSL3-treated populations. Parameters for the fits in (E), (F), (H), (I), and for the basins of attraction if (G) and (J) are come from least-square fitting the mathematical model to the data, see Methods and Document S1.

### Ferroptosis sensitivity is restricted by growth space and not limited RSL3 availability per cell

To control for the possibility that in the LD condition individual cells receive relatively higher doses of RSL3 leading to increased induction of ferroptosis, we repeated RSL3 treatments of cells seeded at LD, MD and HD comparing growth in 24 well versus 12 well plates while scaling the medium volumes and RSL3 amounts to the smaller growth area provided by 24-well plates, thereby keeping RSL3 concentration per cell constant. By doing this, only the growth space, i.e. relative confluency of any given cell number varied between the 12- and 24-well condition. Cells grown in 24-well plates at all three starting densities showed a smaller induction of general ROS upon treatment with RSL3 (Figure 4A) in comparison to the 12-well conditions (Figure 3B). Importantly, the MD condition seeded in 24-well plates was much more resistant to lipid ROS and cell death (Figure 4B, 4C) than the same cell number treated with the same drug concentration per cell in a 12-well format with more growth space (Figure 3C, 3D). Moreover, cultivation under restricted space (24-well) led to unperturbed continued growth of MD cells while confluency of MD cells was decreased under RSL3 treatment with unrestricted space (Figure 4C). These data demonstrate that it is indeed high cellular confluence and not limited drug availability per cell that causes increased ferroptosis resistance and decreased levels of ROS and lipid ROS. In our mathematical model this mechanism emerges from the density-dependent rate of cell divisions, and a rate of ROS production that is proportional to the rate of cell divisions.

**Figure 4.**
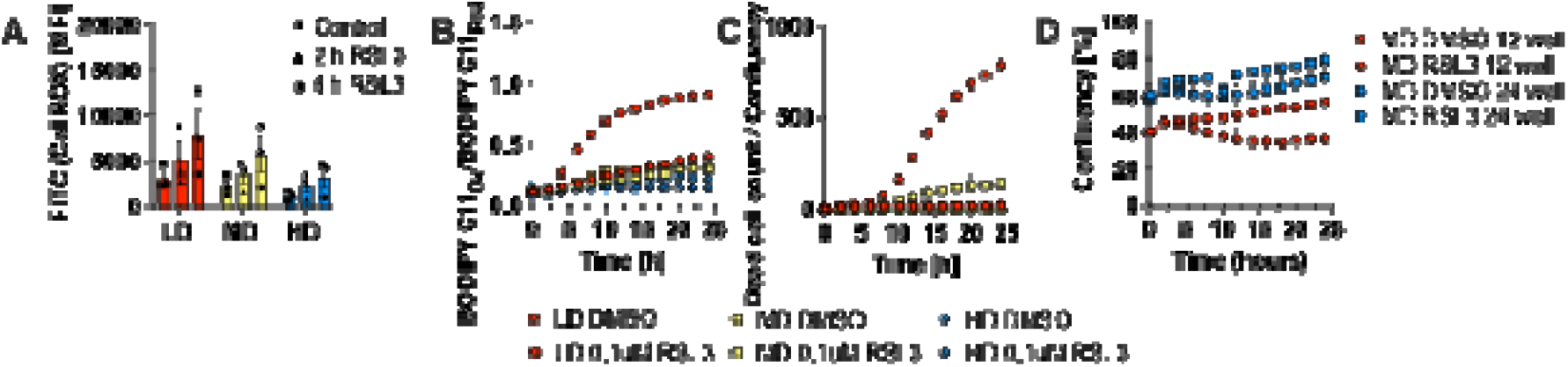
Ferroptosis sensitivity is restricted by growth space and not limited RSL3 availability per cell. (A) Total cellular ROS measured using the CellROX dye in FACS at the indicated time points of MEF populations seeded at LD, MD and HD in 24-well plates in presence of DMSO or 0.1 µM RSL3. (B) Lipid ROS over time shown as oxidized-to-reduced ratio of the STY-BODIPY dye, measured in the Incucyte live-cell imaging system at the indicated time points of MEF populations seeded at LD, MD and HD in 24-well plates presence of DMSO or 0.1 µM RSL3. (C) Dead cell counts over time shown as dead cell count-to-confluence ratio using the DRAQ7 dead cell dye, measured in the Incucyte live-cell imaging system at the indicated time points of MEF populations seeded at LD, MD and HD in 24-well plates in presence of DMSO or 0.1 µM RSL3. (D) Confluency over time of the Incucyte measurements at the indicated time points of MEF populations seeded at MD in 12-well and 24-well plates in presence of DMSO or 0.1 µM RSL3. Confluency data in 12-well plates is taken from the measurements displayed in figure 3 (D). All data are shown as mean ± SEM.

### Ferroptosis sensitivity of growing populations are determined by OXPHOS-derived ROS levels

Next, we aimed to experimentally increase the rate of ROS accumulation per cell division. To this end, we made use of the principle that cells generating energy equivalents in the form of ATP can do so via OXPHOS or through glycolysis(Greene et al., 2022). While the former process is more efficient in generating ATP, this efficiency comes at the cost of electron leakage from the respiratory chain in the mitochondria leading to a high rate of ROS generation(Nolfi-Donegan et al., 2020). Experimentally, cells can be forced to predominantly use OXPHOS for ATP generation by replacing glucose with galactose in growth media leading to increased ROS which should be uncoupled from cell confluency(Aguer et al., 2011). To achieve this, MEF culture media were selectively depleted and repleted for glucose or galactose in glucose-free medium supplemented with insulin-transferrin-selenium (ITS+) instead of calve serum as it is free of glucose and. Notably, ITS+ supplementation is vital for ferroptosis to proceed as selenium is rate-limiting for GPX4 translation and transferrin allows for uptake of iron and is thereby equally crucial for ferroptosis(Gao et al., 2015). Indeed, growing cells in glucose-free medium replenished with galactose but not glucose led to slightly increased ROS in cells grown at HD condition and increased lipid ROS in all three starting densities in comparison to glucose supplemented cells within 24 hours (Figure 5A, 5B). In line with our model, the increase in lipid ROS in the galactose condition also led to increased lipid ROS under RSL3 treatment, resulting in strong sensitization to ferroptotic cell death in galactose-compared to glucose-treated cells at HD (Figure 5C-5E). These data together with our data testing cellular growth space indicate that the inhibitory effect of high population density on ferroptosis cell fate is caused by low metabolic levels of cellular ROS and not cellular proximity.

**Figure 5.**
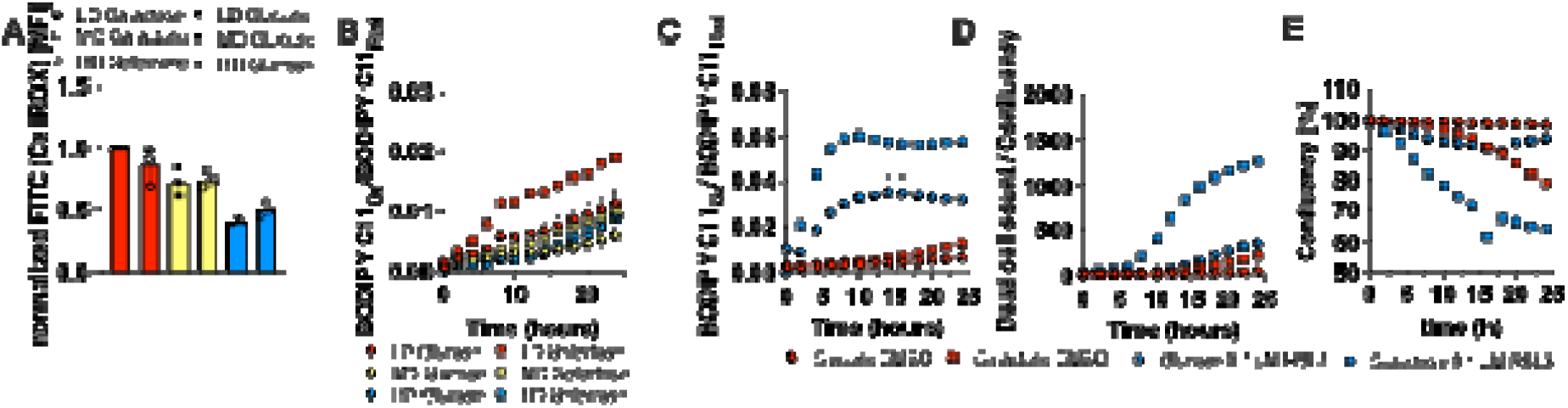
Ferroptosis sensitivity of growing populations are determined by OXPHOS-derived ROS levels. (A) Total cellular ROS measured using the CellROX dye in FACS at the indicated time points of MEF populations seeded at LD, MD and HD in galactose or glucose medium and in presence of DMSO or 0.1 µM RSL3. (B) Lipid ROS over time shown as oxidized-to-reduced ratio of the STY-BODIPY dye, measured in the Incucyte live-cell imaging system at the indicated time points of MEF populations seeded at LD, MD and HD in galactose or glucose medium (C) Lipid ROS over time shown as oxidized-to-reduced ratio of the STY-BODIPY dye, measured in the Incucyte live-cell imaging system at the indicated time points and in presence of DMSO or 0.1 µM RSL3 at HD. (D) Dead cell counts over time shown as dead cell count-to-confluence ratio using the DRAQ7 dead cell dye, measured in the Incucyte live-cell imaging system at the indicated time points of MEF populations seeded at HD in glucose or galactose medium in presence of DMSO or 0.1 µM RSL3. (E) Confluency over time data from measurements displayed in (D). All data are shown as mean ± SEM.

## Discussion

By integrating mathematical modeling with experimental data, we provide evidence that cellular population dynamics, metabolic ROS production, and ferroptosis sensitivity are interlinked. These results have important implications for the design and control of in vitro studies testing ferroptosis sensitivity in fast-growing cell populations such as cancer cell lines. Specifically, we show that low-density (LD) cultures exhibit faster proliferation, elevated ROS generation and greater sensitivity to ferroptosis. This inverse relationship between cellular density and sensitivity to ROS was first established by a study in human fibroblasts(Kim et al., 2012). While the concept of ferroptosis was unknown at the time, more recent studies have provided evidence for a direct link between cellular density and ferroptosis sensitivity. A specific cell state in glioma in which constitutive active NOTCH signaling increases mitochondrial ROS and ferroptosis sensitivity has been reported(Banu et al., 2024). Interestingly, epithelial cancer cell lines were shown to restrict ferroptosis via E-Cadherin-mediated YAP1 sequestration at high cellular densities and cell-to-cell contact(Wu et al., 2019). Moreover, low cell density was shown to render breast cancer cell lines ferroptosis sensitive due to elevated PUFA integration into their lipidomes(Panzilius et al., 2019).

Furthermore, the use of galactose to force mitochondrial oxidative phosphorylation (OXPHOS)(Aguer et al., 2011)—a known source of ROS(Labuschagne et al., 2019)— revealed that metabolic context, rather than density alone, is a key determinant of ferroptosis susceptibility. Interestingly, oxygen consumption in cell populations has long been known to decrease markedly upon reaching confluency(Bereiter-Hahn et al., 1998; Jorjani and Ozturk, 1999). More recent studies have shown that mitochondrial activity decreases with cellular density in epithelial cells(Thurakkal et al., 2023), while oxygen consumption in 3D tissue models using HepG2 cells is markedly increased at lower densities(Magliaro et al., 2019), and proliferating fibroblasts show an increase of OXPHOS by 81% in comparison to quiescent fibroblasts(Yao et al., 2019), a cell state that is commonly induced by high density conditions(Fan and Meyer, 2021). Taken together with our findings, this suggests that the increase in ferroptosis sensitivity in low density and galactose supplemented cell populations observed in this study share upregulation of OXPHOS as a common underlying mechanism. Of note, the elicitation of OXPHOS in proliferating fibroblast population is driven by mitochondrial fusion(Yao et al., 2019), a process by which cells adapt to metabolic challenges such as fasting(Lee et al., 2014), whereas quiescent cells show signs of mitochondrial fragmentation(Yao et al., 2019). In light of our findings linking density-dependent proliferation to OXPHOS and thereby ferroptosis sensitivity, this supports and refines earlier studies indicating that mitochondrial ROS contribute to lipid peroxidation and ferroptosis in specific contexts (Gaschler et al., 2018; Nolfi-Donegan et al., 2020). Our findings extend this concept by showing that seeding density is actively linked with ferroptosis sensitivity through a negative feedback mechanism involving ROS dynamics and proliferation rates. Importantly, our mathematical modeling aligns with the emerging understanding of ferroptosis regulation as a systems-level process with multiple feedback loops(Co et al., 2024). While prior models have considered the role of iron, lipid metabolism, and antioxidant systems(Konstorum et al., 2020), our model uniquely incorporates population growth dynamics, revealing bistable outcomes (ferroptosis-sensitive vs. -resistant states) depending on initial conditions such as cell density and ROS levels. This prediction was experimentally validated using scaled culture formats (12-vs. 24-well plates), ruling out drug dilution as a confounding factor and emphasizing that cell density and growth-induced ROS accumulation drive the feedback loop. This model explains the seemingly paradoxical observation that low-density cell populations—unrestricted by spatial constraints—can initiate ferroptosis. This phenomenon arises from the accumulation of reactive oxygen species (ROS) generated during rapid expansion. In principle, such a mechanism could affect any context of accelerated cellular growth, where instead of establishing a large population, the cells undergo population collapse due to ferroptotic cell death. Central to our model is a negative feedback loop: rapid cell divisions lead to increased ROS production, which induces lipid peroxidation and ultimately triggers ferroptosis. This dynamic gives rise to bistable behavior, with one state marked by low proliferation, minimal ROS levels, and absence of ferroptosis, and the other characterized by high proliferation and turnover, elevated ROS levels, and ferroptotic cell death.

Together, our findings position cell density as both a predictor and regulator of ferroptosis susceptibility, mediated through ROS and metabolic activity. This has broad implications for tissue homeostasis, cancer biology, and therapy resistance, particularly in light of recent interest in exploiting ferroptosis as a therapeutic vulnerability in proliferative and therapy-resistant tumor populations(Hassannia et al., 2019). By integrating experimental data with predictive modeling, our study adds a quantitative framework to the qualitative observations previously reported, offering a deeper mechanistic understanding of how proliferative states and metabolic stress shape ferroptotic outcomes.

## Materials and Methods

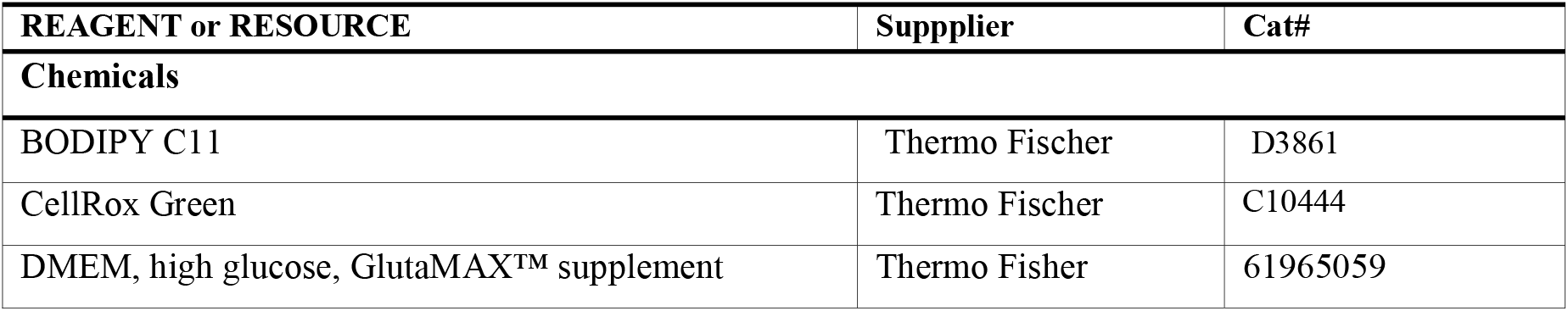
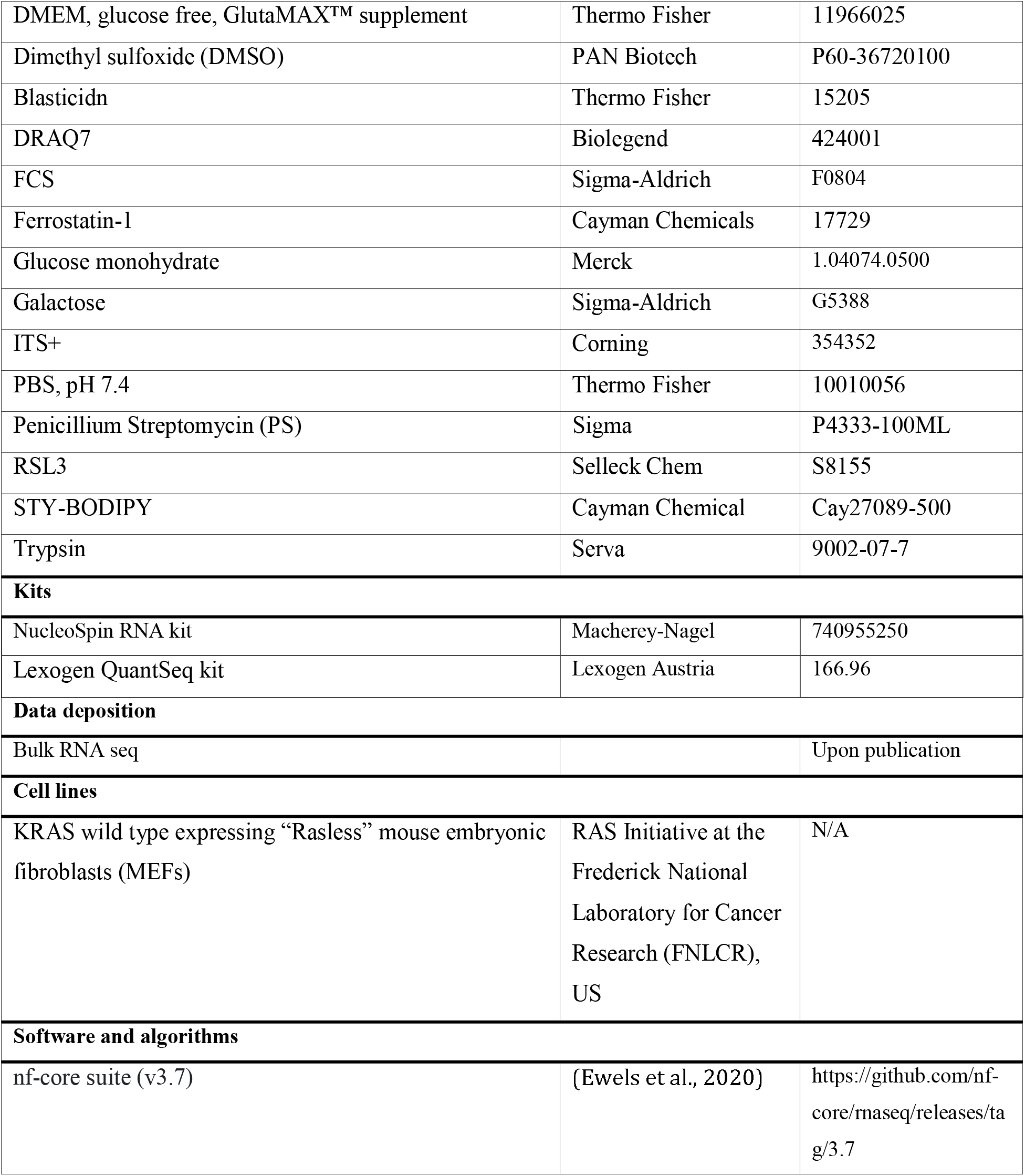
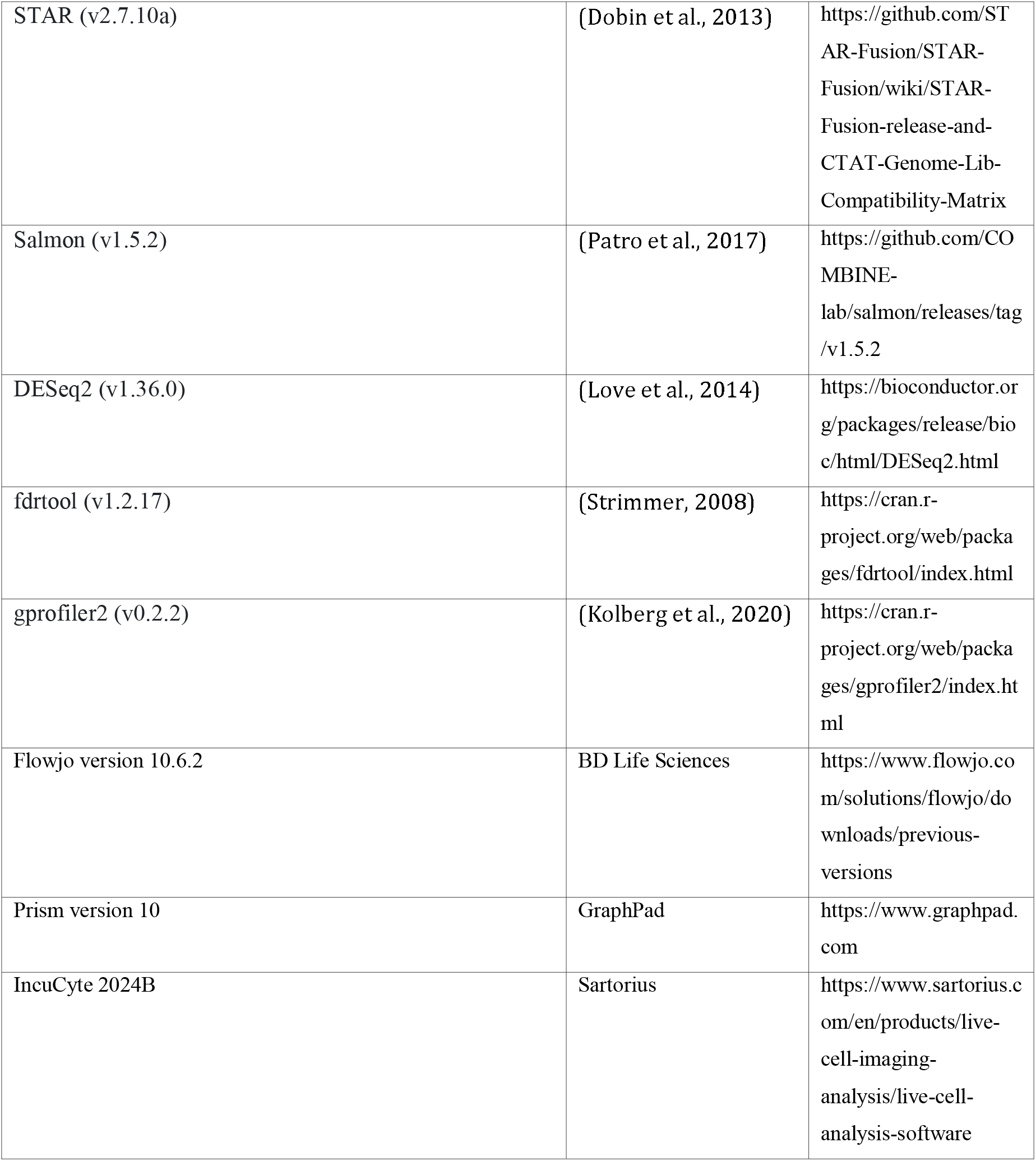

### Cells

Mefs were kept in T-75 flasks at 37°C with 5% CO2 and all media were supplemented with 4 µg/mL blasticidin, 10% FCS and 1% P/S. MEFs were tested for mycoplasma at regular intervals (mycoplasma barcodes, Eurofins Genomics). Cells were passaged every 3-4 days and discarded after 3 weeks.

### Quantification and statistical analysis

Raw data from bulk RNA sequencing was processed as described above, and all other raw data was processed using Excel and GraphPad prism. For statistical testing and generation of figures, GraphPad Prism 10 was used. Two-tailed t-tests were used to compare two conditions, and two-way ANOVA was used to compare multiple samples. All measurements were performed at least three times, and results are presented as mean ± standard error mean (SEM). ns: not significant; ⍰p < 0.05; ⍰⍰, p < 0.01; ⍰⍰⍰, p < 0.001; ⍰⍰⍰⍰, p < 0.0001.

### Population doublings

Cells were seeded in T25-flasks (LD: 329.000; MD: 724.000; HD: 1.578.000 cells per flask) in 5 mL medium and incubated for 24, 48 and 72 h. Afterwards, cells were detached in 1 mL trypsin+ 5mL medium as described above and counted using a Casy Model TT cell counter (Roche Diagnostics). Population doublings were then calculated using the following equation:

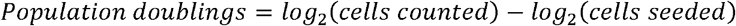

### Reagents

RSL3 was dissolved in DMSO at 1 mM.

### Treatments

For treatment with RSL3, cells were seeded in 24-well plates at 25.000 (low density; LD), 55.000 (medium density; MD) or 120.000 (high density; HD) per well. After 24 hours, RSL3 was added to the medium at a final concentration of 0.1 µM. For glucose/galactose exchange experiments, cells were seeded in DMEM GlutaMAX™ supplemented with blasticidin, PS and FCS as described above. After 24 h, cells were washed thrice with PBS (Gibco) and received glucose-free DMEM (Gibco) supplemented with either 4,5 g/L glucose or galactose, 4 µg/mL blasticidin, 1% P/S and 1% Insulin-Selenium-Transferrin+ (ITS+) supplement (Corning).

### RNA sequencing

For RNA sequencing, MEFs were plated in a 6 well plate at 125.000 (LD) or 600.000 (HD) cells per well. After 24 h, cells were washed with PBS and RNA isolation was done using the NucleoSpin RNA kit (Macherey-Nagel). cDNA libraries amplified from the 3⍰ UTR were generated from total RNA using the Lexogen QuantSeq kit (Lexogen, Austria) according to the standard protocol and sequenced with a 50-bp single-end protocol on Illumina HiSeq4000 sequencer (Illumina, USA).

Primary data analysis was conducted using the RNA-seq pipeline from the nf-core suite (v3.7)(Ewels et al., 2020); sequencing reads were aligned to the GRCh38 (v103) human reference genome using STAR (v2.7.10a)(Dobin et al., 2013). Gene quantification was conducted using Salmon (v1.5.2)(Patro et al., 2017). The pipeline was executed with default parameters. Downstream differential expression analysis was performed using DESeq2 (v1.36.0)(Love et al., 2014), with default parameters. To enhance the accuracy of fold-change estimation, we included mouse ID as a batch effect in the design matrix. For some comparison, the original p-values inferred by DESeq2 revealed significant deviation from the expected uniform null distribution, suggesting low sensitivity. To correct for this, we recomputed the raw p-values using fdrtool (v1.2.17)(Strimmer, 2008) to increase the power of the differential expression procedure while maintaining efficient control for false discovery. Subsequently, the Benjamini-Hochberg procedure was applied to correct the p-values for multiple tests.

GO enrichment analysis was conducted using gprofiler2 (v0.2.2)(Kolberg et al., 2020). The selection criteria focused on differentially expressed genes, as defined above. Using ordered gene query and gProfiler’s “g_SCS” method for p-value adjustment, which accounts for the hierarchical structure of GO terms, enriched GO terms were identified among the differentially expressed genes.

### Flow cytometry

MEFs were seeded in 24 or 12 well plates, treated after 24 with RSL3 as described above and incubated for the indicated durations. BODIPY C11 was added to each well at a final concentration of 5 µM followed by incubation for 30 min at 37°C. Cells were washed, detached and the cell pellet was then resuspended in 200 μl of PBS with 2% FCS and [1 *μ*g/ml] propidium iodide (PI). CellROX was added to each well at a final concentration of 2.5 µM followed by incubation for 30 min at 37°C. Cells were washed, detached and the cell pellet was then resuspended in 200 μl of PBS with 2% FCS and [1 *μ*g/ml] propidium iodide (PI). Flow cytometry data were acquired on the BD LSRFortessa (BD Biosciences) and analyzed with Flowjo V10.6.2.

### Live cell imaging (IncuCyte)

MEFs were plated in 12- and 24-well plates and treated as indicated. For dead cell quantification DRAQ7 [100 mM] was used. For lipid ROS quantification STY-BODIPY [1 *μ*M] was used. Cells were imaged every 2h using the 10x objective within the IncuCyte SX5 live cell imaging system (Sartorius). Analysis for confluence, DRAQ7-positive (dead), reduced- and oxidized-BODIPY positive cells was performed using the Software IncuCyte 2024B (Sartorius). To quantify the number of live cells (live cell confluence) and their lipid ROS levels we carried out additional image analysis to estimate the fraction of the Incucyte signal contributed from dead cells and from living cells. We converted all video frames and images per vessel and applied a color mask to exclude dead cells based on their DRAQ7 signal. Exclusion of the signal from dead cells was performed by expanding the dead cell mask and identifying its overlap with the fluorescence and coverage mask, see Document S1.

### Cellular ROS quantification

To quantify the level of cellular lipid ROS (lipid ROS levels averaged over live cells) we normalize the lipid ROS signal from live cells by dividing by the live cell confluence.

### Mathematical model and data fitting

The first differential equation in Fig 2B gives the rate of change of cell numbers relative to 100% confluence (left hand side) as a balance between the rates of cell divisions and cell death, respectively (right hand side). Analogously, the second equation gives the rate of change of lipid ROS levels (left hand side) and lipid ROS generation and clearance rates (right hand side). We estimate the model parameters by fitting the solution of the differential equation in Fig. 2B to the data using a weighted least-square fit. These parameter estimates give the fits to the time courses in Fig. 4 E, F, H, I and the basins of attraction in Fig. 4 G and J. Details are given in Document S1.

## Supporting information

Supplementary Information: Mathematical modelling and data analysis

## Acknowledgments

This work was funded through a collaborative research center grant on predictability in evolution (CRC1310, project ID 325931972, SvK and J. Berg), cell death (CRC1403, project ID 414786233, SvK), small cell lung cancer (CRC1399, project ID 413326622, SvK) and B-cell lymphomas (CRC1530, project ID 455784452., SvK) and a priority program on ferroptosis (SPP2306, project ID 461704389, SvK), all funded by the German Research Foundation (Deutsche Forschungsgemeinschaft, DFG), an eMed consortium grant by the BMBF (InCa-01ZX2201A, SvK), via CANTAR (SvK) which is funded through the program “Netzwerke 2021”, an initiative of the Ministry of Culture and Science of the State of Northrhine Westphalia, Germany and a project grant (A07, SvK) funded by the center for molecular medicine cologne (CMMC) Cologne.

## Author Contributions

ES, EYI, J. Berg and SvK conceived the project and wrote the manuscript. SvK and J. Berg supervised the project. EYI and ES performed the majority of experiments, modelling and analysis with significant contributions from FM, FIY and J. Brägelmann.

## Data Availability

RNAseq datasets generated within this study have been deposited at public repositories (please refer to key resource table) and are publicly available upon publication. All other original data published within this study are available from the lead contact upon request.

The code used for mathematical modelling (see Document S1) will be made available upon publication.

Any additional information required to reanalyse the data reported in this work paper is available from the corresponding author upon request.

## References

Aguer C, Gambarotta D, Mailloux RJ, Moffat C, Dent R, McPherson R, Harper M-E. 2011. Galactose enhances oxidative metabolism and reveals mitochondrial dysfunction in human primary muscle cells. PLoS One 6:e28536. doi:10.1371/journal.pone.0028536

Banu MA, Dovas A, Argenziano MG, Zhao W, Sperring CP, Cuervo Grajal H, Liu Z, Higgins DM, Amini M, Pereira B, Ye LF, Mahajan A, Humala N, Furnari JL, Upadhyayula PS, Zandkarimi F, Nguyen TT, Teasley D, Wu PB, Hai L, Karan C, Dowdy T, Razavilar A, Siegelin MD, Kitajewski J, Larion M, Bruce JN, Stockwell BR, Sims PA, Canoll P. 2024. A cell state-specific metabolic vulnerability to GPX4-dependent ferroptosis in glioblastoma. EMBO J 43:4492–4521. doi:10.1038/s44318-024-00176-4

Bebber CM, Müller F, Clemente LP, Weber J, Karstedt S von. 2020. Ferroptosis in Cancer Cell Biology. Cancers 12:164.

Bereiter-Hahn J, Münnich A, Woiteneck P. 1998. Dependence of Energy Metabolism on the Density of Cells in Culture. Cell Struct Funct 23:85–93. doi:10.1247/csf.23.85

Co HKC, Wu C-C, Lee Y-C, Chen S. 2024. Emergence of large-scale cell death through ferroptotic trigger waves. Nature 631:654–662. doi:10.1038/s41586-024-07623-6

Conrad M, Pratt DA. 2019. The chemical basis of ferroptosis. Nature chemical biology 15:1137–1147.

Dixon SJ, Lemberg KM, Lamprecht MR, Skouta R, Zaitsev EM, Gleason CE, Patel DN, Bauer AJ, Cantley AM, Yang WS, III BM, Stockwell BR. 2012. Ferroptosis: An Iron-Dependent Form of Nonapoptotic Cell Death. Cell 149:1060–1072.

Dobin A, Davis CA, Schlesinger F, Drenkow J, Zaleski C, Jha S, Batut P, Chaisson M, Gingeras TR. 2013. STAR: ultrafast universal RNA-seq aligner. Bioinformatics 29:15–21. doi:10.1093/bioinformatics/bts635

Eagle H. 1960. The sustained growth of human and animal cells in a protein-free environment. Proceedings of the National Academy of Sciences 46:427–432. doi:10.1073/pnas.46.4.427

Ewels PA, Peltzer A, Fillinger S, Patel H, Alneberg J, Wilm A, Garcia MU, Tommaso PD, Nahnsen S. 2020. The nf-core framework for community-curated bioinformatics pipelines. Nature Biotechnology 38:276–278. doi:10.1038/s41587-020-0439-x

Fan Y, Meyer T. 2021. Molecular control of cell density-mediated exit to quiescence. Cell Reports 36. doi:10.1016/j.celrep.2021.109436

Friedmann Angeli JP, Schneider M, Proneth B, Tyurina YY, Tyurin VA, Hammond VJ, Herbach N, Aichler M, Walch A, Eggenhofer E, Basavarajappa D, Rådmark O, Kobayashi S, Seibt T, Beck H, Neff F, Esposito I, Wanke R, Förster H, Yefremova O, Heinrichmeyer M, Bornkamm GW, Geissler EK, Thomas SB, Stockwell BR, O’Donnell VB, Kagan VE, Schick JA, Conrad M. 2014. Inactivation of the ferroptosis regulator Gpx4 triggers acute renal failure in mice. Nat Cell Biol 16:1180–1191. doi:10.1038/ncb3064

Gao M, Monian P, Quadri N, Ramasamy R, Jiang X. 2015. Glutaminolysis and Transferrin Regulate Ferroptosis. Molecular cell 59:298–308.

Gao M, Yi J, Zhu J, Minikes AM, Monian P, Thompson CB, Jiang X. 2018. Role of Mitochondria in Ferroptosis. Molecular cell.

Greene J, Segaran A, Lord S. 2022. Targeting OXPHOS and the electron transport chain in cancer; Molecular and therapeutic implications. Seminars in Cancer Biology 86:851–859. doi:10.1016/j.semcancer.2022.02.002

Hassannia B, Vandenabeele P, Berghe TV. 2019. Targeting Ferroptosis to Iron Out Cancer. Cancer cell 35:830–849.

Homma T, Kobayashi S, Sato H, Fujii J. 2021. Superoxide produced by mitochondrial complex III plays a pivotal role in the execution of ferroptosis induced by cysteine starvation. Arch Biochem Biophys 700:108775. doi:10.1016/j.abb.2021.108775

Ivanova JS, Pugovkina NA, Neganova IE, Kozhukharova IV, Nikolsky NN, Lyublinskaya OG. 2021. Cell cycle-coupled changes in the level of reactive oxygen species support the proliferation of human pluripotent stem cells. STEM CELLS 39:1671–1687. doi:10.1002/stem.3450

Jorjani P, Ozturk SS. 1999. Effects of cell density and temperature on oxygen consumption rate for different mammalian cell lines. Biotechnol Bioeng 64:349–356. doi:10.1002/(sici)1097-0290(19990805)64:3<349::aid-bit11>3.0.co;2-v

Kagan VE, Mao G, Qu F, Angeli JPF, Doll S, Croix CS, Dar HH, Liu B, Tyurin VA, Ritov VB, Kapralov AA, Amoscato AA, Jiang J, Anthonymuthu T, Mohammadyani D, Yang Q, Proneth B, Klein-Seetharaman J, Watkins S, Bahar I, Greenberger J, Mallampalli RK, Stockwell BR, Tyurina YY, Conrad M, Bayır H. 2017. Oxidized arachidonic and adrenic PEs navigate cells to ferroptosis. Nature chemical biology 13:81–90.

Kim DP, Yahav J, Sperandeo M, Maloney L, McTigue M, Lin F, Clark RAF. 2012. High cell density attenuates reactive oxygen species: Implications for in vitro assays. Wound Repair and Regeneration 20:74–82. doi:10.1111/j.1524-475X.2011.00746.x

Kolberg L, Raudvere U, Kuzmin I, Vilo J, Peterson H. 2020. gprofiler2 – an R package for gene list functional enrichment analysis and namespace conversion toolset g:Profiler. F1000Research 9:ELIXIR-709. doi:10.12688/f1000research.24956.2

Konstorum A, Tesfay L, Paul BT, Torti FM, Laubenbacher RC, Torti SV. 2020. Systems biology of ferroptosis: A modeling approach. J Theor Biol 493:110222. doi:10.1016/j.jtbi.2020.110222

Labuschagne CF, Cheung EC, Blagih J, Domart M-C, Vousden KH. 2019. Cell Clustering Promotes a Metabolic Switch that Supports Metastatic Colonization. Cell Metabolism 30:720-734.e5. doi:10.1016/j.cmet.2019.07.014

Lee J-Y, Kapur M, Li M, Choi M-C, Choi S, Kim H-J, Kim I, Lee E, Taylor JP, Yao T-P. 2014. MFN1 deacetylation activates adaptive mitochondrial fusion and protects metabolically challenged mitochondria. J Cell Sci 127:4954–4963. doi:10.1242/jcs.157321

Love MI, Huber W, Anders S. 2014. Moderated estimation of fold change and dispersion for RNA-seq data with DESeq2. Genome Biology 15:550. doi:10.1186/s13059-014-0550-8

Magliaro C, Mattei G, Iacoangeli F, Corti A, Piemonte V, Ahluwalia A. 2019. Oxygen Consumption Characteristics in 3D Constructs Depend on Cell Density. Front Bioeng Biotechnol 7:251. doi:10.3389/fbioe.2019.00251

Martens GA, Cai Y, Hinke S, Stangé G, Casteele MV de, Pipeleers D. 2005. Glucose Suppresses Superoxide Generation in Metabolically Responsive Pancreatic β Cells *. Journal of Biological Chemistry 280:20389–20396. doi:10.1074/jbc.M411869200

Nolfi-Donegan D, Braganza A, Shiva S. 2020. Mitochondrial electron transport chain: Oxidative phosphorylation, oxidant production, and methods of measurement. Redox Biology 37:101674. doi:10.1016/j.redox.2020.101674

Panzilius E, Holstein F, Dehairs J, Planque M, Toerne C von, Koenig A-C, Doll S, Bannier-Hélaouët M, Ganz HM, Hauck SM, Talebi A, Swinnen JV, Fendt S-M, Angeli JPF, Conrad M, Scheel CH. 2019. Cell density-dependent ferroptosis in breast cancer is induced by accumulation of polyunsaturated fatty acid-enriched triacylglycerides. doi:10.1101/417949

Patro R, Duggal G, Love MI, Irizarry RA, Kingsford C. 2017. Salmon provides fast and bias-aware quantification of transcript expression. Nature Methods 14:417–419. doi:10.1038/nmeth.4197

Pedrera L, Espiritu RA, Ros U, Weber J, Schmitt A, Stroh J, Hailfinger S, Karstedt S von, García-Sáez AJ. 2020. Ferroptotic pores induce Ca 2+ fluxes and ESCRT-III activation to modulate cell death kinetics. Cell death and differentiation 149:1–14.

Riegman M, Sagie L, Galed C, Levin T, Steinberg N, Dixon SJ, Wiesner U, Bradbury MS, Niethammer P, Zaritsky A, Overholtzer M. 2020. Ferroptosis occurs through an osmotic mechanism and propagates independently of cell rupture. Nat Cell Biol 22:1042–1048. doi:10.1038/s41556-020-0565-1

Schneider M, Wortmann M, Mandal PK, Arpornchayanon W, Jannasch K, Alves F, Strieth S, Conrad M, Beck H. 2010. Absence of glutathione peroxidase 4 affects tumor angiogenesis through increased 12/15-lipoxygenase activity. Neoplasia 12:254–263. doi:10.1593/neo.91782

Seiler A, Schneider M, Förster H, Roth S, Wirth EK, Culmsee C, Plesnila N, Kremmer E, Rådmark O, Wurst W, Bornkamm GW, Schweizer U, Conrad M. 2008. Glutathione peroxidase 4 senses and translates oxidative stress into 12/15-lipoxygenase dependent-and AIF-mediated cell death. Cell metabolism 8:237–248.

Strimmer K. 2008. A unified approach to false discovery rate estimation. BMC Bioinformatics 9:303. doi:10.1186/1471-2105-9-303

Thurakkal B, Hari K, Marwaha R, Karki S, Jolly MK, Das T. 2023. Collective heterogeneity of mitochondrial potential in contact inhibition of proliferation. Biophysical Journal 122:3909–3923. doi:10.1016/j.bpj.2023.08.014

Vucetic M, Daher B, Cassim S, Meira W, Pouyssegur J. 2020. Together we stand, apart we fall: how cell-to-cell contact/interplay provides resistance to ferroptosis. Cell Death Dis 11:789. doi:10.1038/s41419-020-02994-w

Wiernicki B, Dubois H, Tyurina YY, Hassannia B, Bayir H, Kagan VE, Vandenabeele P, Wullaert A, Berghe TV. 2020. Excessive phospholipid peroxidation distinguishes ferroptosis from other cell death modes including pyroptosis. Cell Death and Disease 11:922–11.

Wu J, Minikes AM, Gao M, Bian H, Li Y, Stockwell BR, Chen Z-N, Jiang X. 2019. Intercellular interaction dictates cancer cell ferroptosis via NF2-YAP signalling. Nature 171:273.

Yan H, Tuo Q, Lei P. 2024. Cell density impacts the susceptibility to ferroptosis by modulating IRP1-mediated iron homeostasis. Journal of Neurochemistry 168:1359– 1373. doi:10.1111/jnc.16085

Yang WS, SriRamaratnam R, Welsch ME, Shimada K, Skouta R, Viswanathan VS, Cheah JH, Clemons PA, Shamji AF, Clish CB, Brown LM, Girotti AW, Cornish VW, Schreiber SL, Stockwell BR. 2014. Regulation of Ferroptotic Cancer Cell Death by GPX4. Cell 156:317–331.

Yang WS, Stockwell BR. 2008. Synthetic Lethal Screening Identifies Compounds Activating Iron-Dependent, Nonapoptotic Cell Death in Oncogenic-RAS-Harboring Cancer Cells. Chemistry & Biology 15:234–245.

Yao C-H, Wang R, Wang Y, Kung C-P, Weber JD, Patti GJ. 2019. Mitochondrial fusion supports increased oxidative phosphorylation during cell proliferation. eLife 8:e41351. doi:10.7554/eLife.41351

