## Supplementary Information: Mathematical modelling and data analysis for "An integrated model of population growth saturation and basal ROS levels predicts cellular ferroptosis sensitivity"

**4 Addendum: Fixed points without positive feedback 11**

### 1 A mathematical model linking population growth and lipid ROS dynamics

In the following we set up a quantitative model describing the feedback between population growth, lipid ROS levels, and ferroptotic cell death. At its core is a system of bistable differential equations based on the production of lipid ROS as a side product of cell growth and ferroptotic cell death triggered by lipid peroxidation products (LPO) produced by lipid ROS. We show how the model explains how ferroptosis sensitivity depends on growth rate and initial cell density. Furthermore, we link this model with experimental data by fitting the model parameters to image-based measurements of populations with different initial conditions, from which we derive system parameters and an underlying phase space. In particular, we investigate perturbations of system parameters and explain how decreasing the LPO clearance can push a population towards ferroptotic cell death.

Our mathematical model links population growth, reactive oxygen species (lipid ROS) levels, and ferroptotic cell death. It consists of three mechanisms: 1) A logistically growing population, 2) lipid ROS emerging as a by-product of fast growth and producing lipid ROS, and 3) lipid ROS generating lipid peroxidation products (LPO) which can trigger ferroptotic cell death. We focus on the population size and levels of lipid ROS as key variables of the model. In order to make the model as simple as possible we do not explicitly model the intermediate step of lipid ROS production en route to lipid ROS, or lipid peroxidation by lipid ROS.

Our starting point is the standard logistic growth of a population where cells divide at a rate $a(1 - \frac{n}{K})$. Hereby, $a$ is the basal rate of cell division per cell, $n$ the living population size, and $K$ the carrying capacity. The growth rate thus slows down as the carrying capacity of the population is reached. High levels of lipid ROS trigger ferroptotic cell death (via production of LPO), which we model by a Hill-curve for the rate at which individual cells die $b\frac{r^h}{r^h+r_0^h}$ depending on the level of lipid ROS per cell $r$. $b$ is the maximum rate of cell death and the death rate rises when a lipid ROS level threshold $r_0$ is reached. $r_0$ represents the lipid ROS concentration that causes a half-maximum cell death response. The Hill-coefficient $h$ quantifies the sharpness of that response. The balance between cell birth and death gives a dynamic equation for the population size

$$\frac{dn(t)}{dt} = a\left(1 - \frac{n(t)}{K}\right)n(t) - b\frac{r(t)^h}{r(t)^h + r_0^h}n(t). \tag{1}$$

The second component of the model is a dynamical equation for the level of lipid ROS. ROS emerges as a side product of fast growth and produces lipid ROS. We model this effective process by a lipid ROS level (averaged over cells) which increases by $\alpha$ per cell division. Additionally, we incorporate a lipid ROS-induced positive feedback loop in the lipid ROS accumulation. High levels of lipid ROS can trigger a self-amplified production of more lipid ROS.[1] As in Co et. al,[1] this is modelled by a Hill-function for the rate at which lipid ROS is produced $f\frac{r^l}{r^l+r_1^l}$. $f$ represents a basal feedback rate and $l$ and $r_1$ define the sharpness of the response curve and the half-maximum response lipid ROS concentration. Lipid ROS levels in a cell decrease at a constant rate $\beta$ per lipid ROS molecule due to lipid ROS scavenging and degradation. The balance between lipid ROS generation and degradtion then gives

$$\frac{dr(t)}{dt} = \alpha a\left(1 - \frac{n(t)}{K}\right) + f\frac{r(t)^l}{r(t)^l + r_1^l} - \beta r(t). \tag{2}$$

Equations (1) and (2) specify a deterministic model for the coupled dynamics of the population size and the average level of lipid ROS per cell. It describes a negative feedback loop, where rapid growth leads to high levels of lipid ROS, which lead to ferroptotic cell death, and thus a decrease in the population size.

An attractor basin plot shows the dynamics of this model under different initial conditions. Figure 1A shows population size $n$ and lipid ROS level $r$ on the two axes, and arrows visualise the dynamics defined by equations (1) and (2). One sees that starting from low population sizes and high levels of lipid ROS, a population is taken to an asymptotic state with high rate of cell death and low population size. This ferroptosis-sensitive state is indicated by a red point. On the other hand, initial conditions with high population size and low levels of lipid ROS evolve to an asymptotic state characterised by a low rate of ferroptotic cell death and a population reaching carrying capacity. This ferroptosis-resistant state is indicated by the blue point. The basins of attraction of these asymptotic states are defined by initial conditions that end up in the susceptible or resistant state. These points are shown in red and blue, respectively. The property of the system to support two long-term fates of the population, which are selected by initial conditions, is called bistability.

Figure 1A also shows three representative trajectories, starting at low, medium, and high cell densities and low lipid ROS. For the trajectories starting at medium and high density (shown in yellow and blue respectively), growth is slow as the populations are near the carrying capacity. Hence, the rate of lipid ROS accumulation is low, ferroptosis is not triggered and the population eventually reaches the ferroptosis-resistant state. A population starting at at low cell density (shown in red) initially grows quickly and therefore accumulates higher levels of lipid ROS, triggering cell death. The population does not grow out to the carrying capacity and asymptotically ends up in the ferroptosis-susceptible state at low cell densities. Figure 1B shows the population growth over time for these very representative trajectories, Figure 1C shows the corresponding levels of lipid ROS over time.

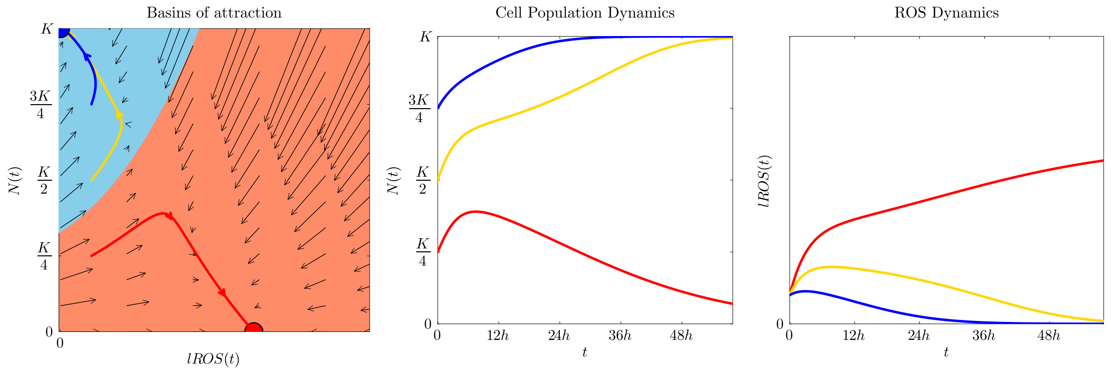

Figure 1: **Bistability and sensitivity to initial conditions predicted by model**. (A) Arrows in this diagram show how population sizes and lipid ROS levels change over time under the dynamics defined by (1) and (2). Colours indicate which asymptotic state a particular point will eventually be taken to; points in the orange area will be taken to the point marked in red (ferroptosis sensitive, high lipid ROS level and low population size), points in the blue area will grow to confluence and have low levels of lipid ROS (point marked in blue). Three representative trajectories, starting at low, medium and high cell densities are shown in red, yellow, and blue. (B) For these three trajectories, the population size is shown over time. (C) Analogously, lipid ROS levels over time are shown. Parameters chosen: $f = 0$, $h = 2$, $a = 0.25$, $b = 0.5$, $K = 1$, $r_0 = 1$ $\alpha = 1$, $\beta = 0.2$

##### 1.1 Model parameters

The bistability of the system and the resulting sensitivity to the initial conditions depends on the choice of parameters. The stability or even the existence of fixed points can change when those parameters are changed. The parameters of this model can be manipulated experimentally. For instance inhibiting LPO reduction mechanisms, for instance by inhibition of GPX4, increases the rate at which LPO accumulate. This means the LPO threshold at which ferroptosis is triggered is reached already at lower levels of lipid ROS. Hence inhibiting LPO reduction is modelled by a decrease in the threshold $r_0$ in the dynamical equation for population size $n$, equation (1). The basal lipid ROS production rate $\alpha$ can be increased for instance by switching from glucose to galactose. Replacing glucose by galactose in cellular growth media forces cells to predominantly use oxidative phosphorylation instead of glycolysis for ATP production, which in turn leads to an increased lipid ROS production. The lipid ROS clearing rate $\beta$ can be reduced by inhibiting the antioxidant defense (ROS scavengers).

Parameter changes of this kind can cause steady states and the basin boundary to shift. As a result, populations with very similar initial conditions can be directed to different states by changing the model parameters. A population that survives under one set of parameters can be pushed towards extinction if a critical parameter is changed.

#### 2 Data analysis

##### 2.1 Experimental design and data

We performed experiments with mouse embryonic fibroblasts (MEFs) seeded at low (LD), middle (MD) and high cell density (HD) into cell culture wells each with 25,000, 55,000 and 120,000 cells, respectively. The experiments were realised in 12-well plates. In this section, we describe in detail we analyzed the resulting data, in the following sections we discuss the inference of model parameters from this data. For reasons discussed below, we do not simply use aggregate fluorescence signals, but use a preprocessing step based on a frame-by-frame analysis of individual cells.

For every condition cell confluence, dead cells and lipid ROS were monitored every 2 hours over a 24-hour period using live cell imaging (Incucyte™). The level of lipid ROS was measured using the fluorescent lipid ROS dye BODIPY™and occurrence of cell death was measured using the fluorescent dye Draq7™. For each density (LD, MD, HD), three replicates were performed in a control medium with glucose (Ctrl) and, for comparison, three replicates in which the MEFs were additionally treated. Therefore, MEFs were seeded in LD, MD and HD and left to grow in the presence or absence of RSL3 to observe the main differences in cell survival and compare the causes.

The data presented in Figure 2 show the following: Cells under control conditions grow towards confluence independent of the seeding density and the cell death rate is low. When the population is treated with RSL3, the growth rate and death rate remains unchanged at high cell densities. High cell confluence thereby prevents cell death and populations do not undergo ferroptosis. MEFs seeded in LD, on the other hand, show the highest lipid ROS levels under RSL3 and fail to grow to confluence. Populations at low cell densities therefore show ferroptosis susceptibility and go extinct. The cell death rate is elevated. The experimental data shows bistability and sensitivity to initial conditions, with population density being the main predictor of survival or extinction. Changing the experimental conditions can change the long-term fate of a population: by adding the small-molecule inhibitor RSL3, we inhibited GPX4 activity and we in fact observe an increased lipid ROS level, which is directly related to cell density. While the control conditions do not show such bistability, the populations treated with RSL3 show two possible outcomes depending on the initial conditions of the population. The change in a critical parameter ($r_0$) led to a shift in the basin boundary and cell fate.

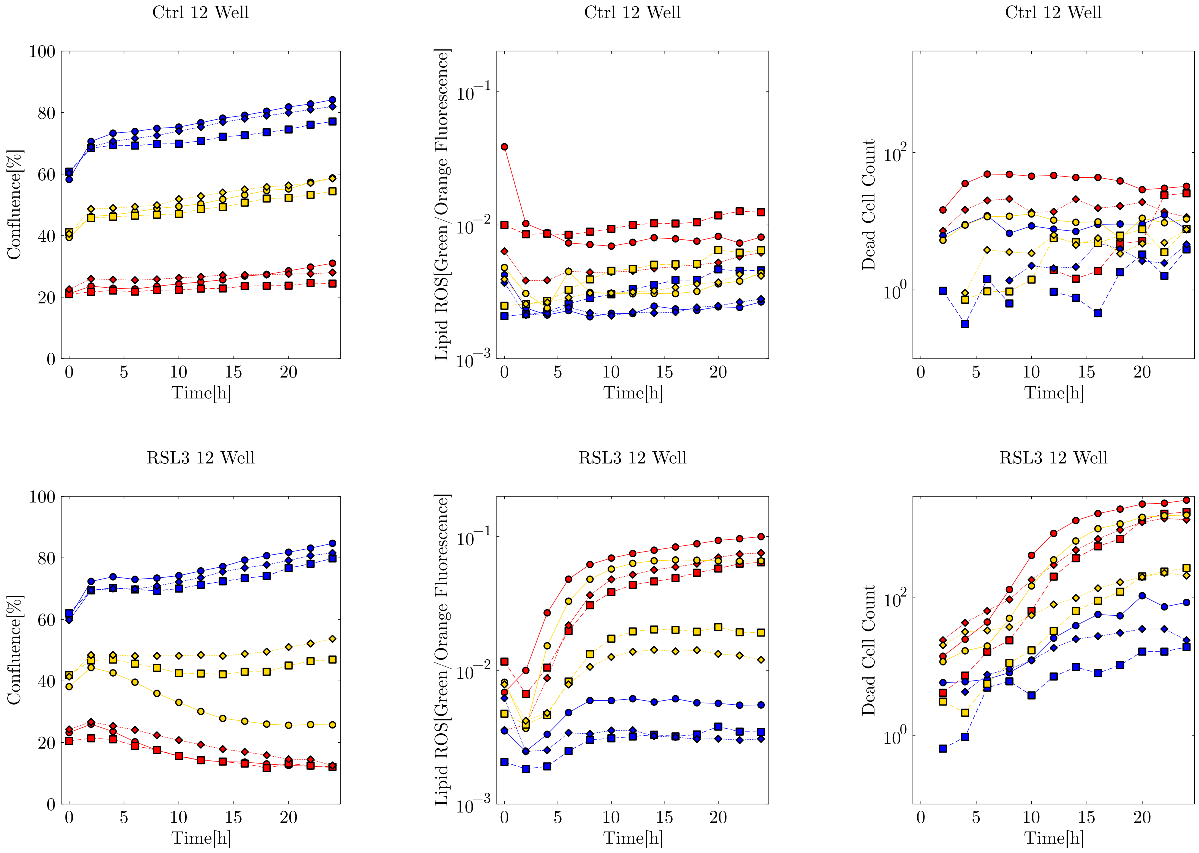

Figure 2: **Comparison of confluence, lipid ROS, and dead cell measurements over time for control and RSL3-treated data**. Three replicates for three initial conditions, starting at low, medium, and high cell densities are shown in red, yellow, and blue. Markers indicate different replicates. The top row presents Ctrl experiments, the bottom row RSL3 experiments.

#### 2.2 Measured observables and mapping to model variables

In the experiments, cell density is quantified using confluence, which represents the percentage of the area of the well covered by cells in each image. This value is proportional to the number of cells and can be mapped to the variable $n$ in our model, which denotes the number of living cells.

Lipid ROS levels are measured through the total fluorescence intensity of a lipid ROS dye. The dye shifts its emission from orange to green upon oxidation, and the integrated intensity of the green signal normalised over the orange signal over all cells is used as a proxy for lipid ROS. We map this fluorescence intensity normalised over the confluence to the variable $r$, which represents the lipid ROS level per cell in the model.

This mapping of the experimental variables to the model variables $n(t)$ and $r(t)$ allows us to interpret the experimental data in terms of the model and enables us to make quantitative model fitting and inference.

#### 2.3 Preprocessing and image analysis

The confluence and lipid ROS fluorescence data also include dead cells, as their remains do not necessarily disappear with cell death. Figure 3 shows an example image that illustrates that both lipid ROS fluorescence and confluence contain contributions from dead cells. Dead cells do not disappear immediately when the cell membrane is compromised and fluorescence becomes visible, and the fluorescent lipid ROS molecules of these cells also persist after cell death. For this reason, we carry out an image analysis to estimate the fraction contributed from dead cells to confluence and lipid ROS measurements and exclude it from the signal. This preprocessing is required to estimate the data from living cells and map it to our model describing the growth and lipid ROS dynamics of a living population.

From video frames we analyse 9 images per vessel by converting each frame from RGB to HSV color space and apply color masks for the lipid ROS and dead cell signal. The lipid ROS signal appears in a yellow-green spectrum, due to oxidation shifting the dye from orange to green. Dead cells appear in blue to cyan color. The color-specific masks are defined by HSV thresholds such that noise is reduced but we still cover typical values and a broad spectrum. For the dead cell staining, we selected hues between 90–130 with moderate to high saturation and brightness (50–255 and 110–255, respectively), which captures typical blue tones while excluding background noise. For the lipid ROS staining, we extended the hue range from 15–60 to cover both greenish-yellow and orange-shifted yellows and we selected moderate to high saturation and brightness (70–255 and 70–255, respectively). This includes less saturated yellow signals but reduces background noise. With this, we compute the total yellow-green fluorescence intensity in an image by summing pixel intensities across the lipid ROS mask. Similarly, cell coverage is estimated via grayscale thresholding and masking. We converted images to grayscale and applied a binary threshold at an intensity of 50. Pixels above this threshold correspond to cell-covered regions.

To then exclude signal originating from dead cells, we expand the dead cell areas outward by applying a morphological dilation to the dead cell mask using a $15 \times 15$ pixel structuring element and identify its overlap with the yellow-green mask or the confluence mask. We make a simplified assumption and use the same dilation of the mask. In this way, we can calculate the fraction of intensity and coverage associated exclusively with dead cells at each time point and in each image per vessel. This serves as a time-dependent quantity to convert the experimental data to more tailored lipid ROS and confluence data attributed only to living cells. The variance of this factor between different images is small and negligible compared to the variance between experimental replicates, thus we use the mean value across all images for preprocessing.

To summarise, our preprocessing pipeline allows us to extract estimates of the density of living cells and the lipid ROS values emerged from living cells over time and interpret them as our model variables $n(t)$ and $r(t)$, respectively. As can already be seen in Figure 2, there are significantly more dead cells in the RSL3 samples than in the Ctrl samples, and therefore these trajectories are also changed more during preprocessing. Figure 4 shows the comparison between the original data and the processed data.

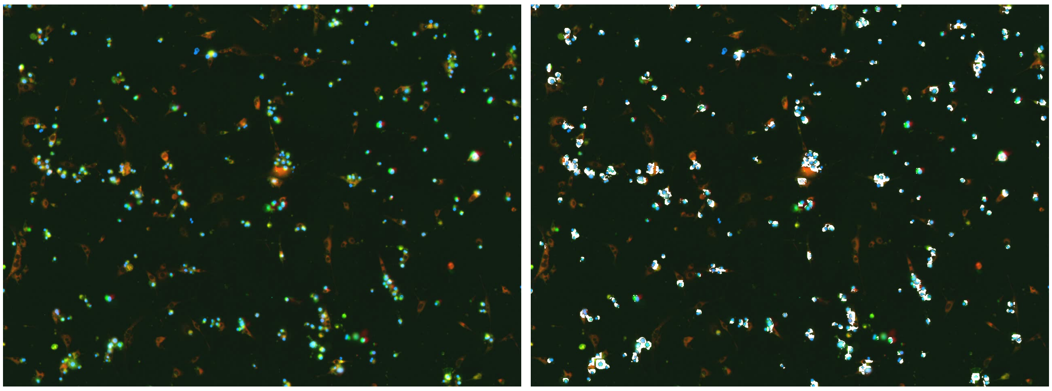

Figure 3: **A representative frame from image based measurements used for pre-processing**. Blue fluorescence marks dead cells, yellow-green fluorescence is lipid ROS. Left: Original frame, Right: White mask illustrating yellow-green signal close to dead cells, which is excluded in the preprocessing. The frame is taken from replicate 1 at MD in a 12-well under RSL3 treatment at $T = 24h$.

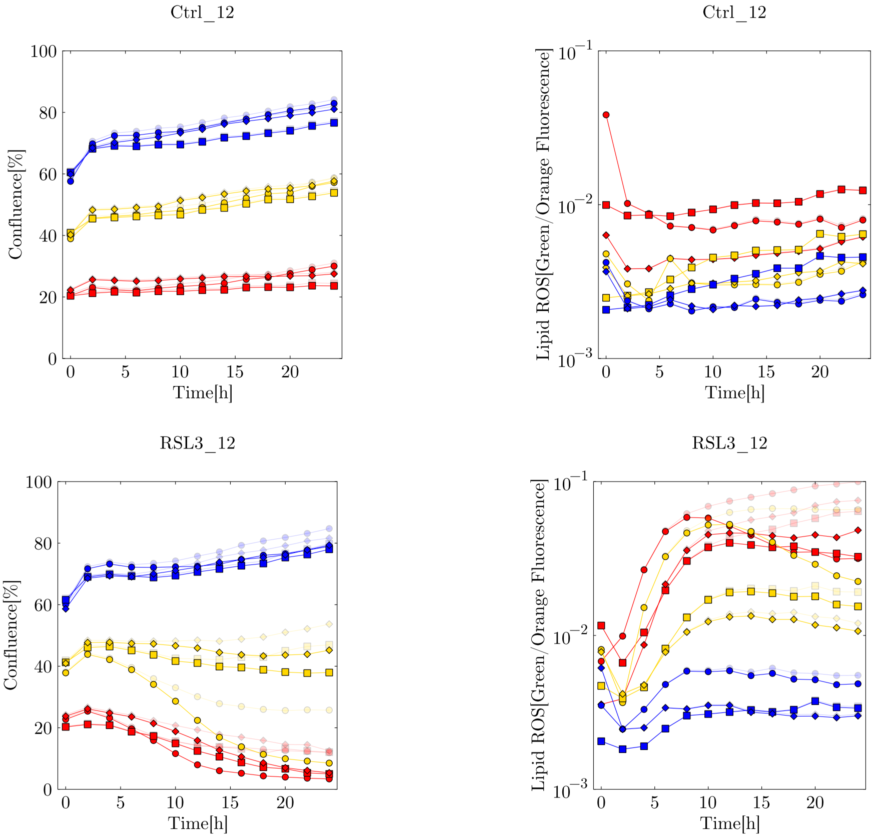

Figure 4: **Comparison of lipid ROS and confluence measurements before vs. after processing via image analysis**. Time series for control (top row) and RSL3-treated data (bottom row). Three replicates for three initial conditions, starting at low, medium and high cell densities are shown in red, yellow, and blue. Markers indicate different replicates. The data before processing is shown with a light line, the data after processing is shown with a bold line.

### 3 Quantitative analysis: fitting model to data

#### 3.1 Parallel fitting to different conditions and treatments

To link the model quantitatively to the experimental data, we perform a simple fit of the model to the data. In this way, we can derive the model parameters from the data. The estimated parameters allow us to compare population growth and lipid ROS rates. The most important part of our fitting scheme is that we fit all time series with initial seeding density LD, MD and HD simultaneously. The crucial aspect is that under one particular condition (with or without RSL3) different populations are seeded and evolve under the same conditions, and are modelled by a common set of parameters. Thus, if populations with different initial conditions reach different asymptotic states, either reaching confluence or not, this must stem from a bistable property system. We even go one step further and assume that only one parameter, namely $r_0$, differs between the control and the treated condition (see Section 1.1). All other parameters are presumed to be the same under both conditions. This assumption is biologically motivated: RSL3 specifically impairs LPO elimination, which primarily affects $r_0$, while the underlying properties of the cells are intrinsic and remain unchanged. Besides that biological motivation, there is also a technical motivation for a joint fit of the control and RSL3 data. In the control condition, lipid ROS accumulation and cell death is low for all seeding densities. This makes it difficult to accurately estimate parameters such as the death rate $b$ or lipid ROS parameters. The absence of cell death and the resulting dynamics can thereby be well explained by multiple parameter combinations of low death rates, high lipid ROS concentration thresholds, or high lipid ROS clearance rates. Hence, the RSL3 condition is richer than the control condition in terms of death-related dynamics. On the other hand, the control data serves as a good reference with minimal cell death to better estimate the cell division rate. For this reason, we simultaneously fit our model to the data from the control and the RSL3-treated populations at different initial densities (LD, MD, HD). We condition most parameters to be equal for all time series to reflect common underlying biological processes, leaving the lipid ROS cell death threshold $r_0$ condition specific. Fit parameters are thus $a, b, \alpha, \beta, f, r_1, r_{0,\mathrm{Ctrl}}, r_{0,\mathrm{RSL3}}$.

#### 3.2 Least-squares fitting

To fit the model to the experimental data, we use a simple least-squares fit. The method of least-squares minimises the sum of the squares of the differences between the observed data and the predicted data. Our weighted least-squares loss function $L(p)$, that quantifies the discrepancy between the model output and data, is defined as

$$L(p) = \sum_{j \in D} \sum_{i=0}^{T} \left[ \frac{\left(n_{\mathrm{data},j}(t_i) - n_{\mathrm{pred},j}(p, t_i)\right)^2}{\sigma^2_{n,\mathrm{data},j}} + \frac{\left(r_{\mathrm{data},j}(t_i) - r_{\mathrm{pred},j}(p, t_i)\right)^2}{\sigma^2_{r,\mathrm{data},j}} \right]. \quad (3)$$

We sum over the data set $D$ and all time points $T$. The experimental data are given in pairs of $(n_{\mathrm{data},j}(t_i), r_{\mathrm{data},j}(t_i))$, while $(n_{\mathrm{pred},j}(p, t_i), r_{\mathrm{pred},j}(p, t_i))$ denote the predicted trajectories obtained from the deterministic equations for replicate $j$ at time $i$ given a parameter guess $p$. The parameter set $p$ contains all variable parameters that are optimised (see section 3.1) as well as the initial conditions $(n_{\mathrm{pred},j}(t = 0), r_{\mathrm{pred},j}(t = 0))$ , which are optimised as well. We define a normalisation using the variance of a time series to balance the different scales of the confluence data $n$ and the lipid ROS data $r$. $\sigma_{n,\mathrm{data},j}$ and $\sigma_{r,\mathrm{data},j}$ represent the variance of trajectory $j$ over a time period $T$.

In this approach, the best-fitting parameter set for the data is the set that minimises the sum of all weighted squared differences of every seeding density (LD, MD, HD) and condition (Ctrl, RSL3): $p_{\mathrm{opt}} = \arg\min_p L(p)$.

##### 3.3 Data used for fits

The set $D$ denotes the collection of all trajectories used for model fitting. Depending on the fitting strategy, $D$ can represent the set of mean time evolutions, i.e., the mean across replicates for each seeding density

$$D_{\text{mean}} = \left\{ \begin{pmatrix} \langle n_{\text{LD, Ctrl}}(t) \rangle \\ \langle r_{\text{LD, Ctrl}}(t) \rangle \end{pmatrix}, \begin{pmatrix} \langle n_{\text{LD, RSL3}}(t) \rangle \\ \langle r_{\text{LD, RSL3}}(t) \rangle \end{pmatrix}, \begin{pmatrix} \langle n_{\text{MD, RSL3}}(t) \rangle \\ \langle r_{\text{MD, RSL3}}(t) \rangle \end{pmatrix}, \begin{pmatrix} \langle n_{\text{HD, Ctrl}}(t) \rangle \\ \langle r_{\text{HD, Ctrl}}(t) \rangle \end{pmatrix}, \dots \right\},$$

or the full set of individual replicates across all experimental conditions in order to capture biological variability

$$D_{\text{rep}} = \left\{ \begin{pmatrix} n_{\text{LD, Ctrl,1}}(t) \\ r_{\text{LD, Ctrl,1}}(t) \end{pmatrix}, \begin{pmatrix} n_{\text{LD, Ctrl,2}}(t) \\ r_{\text{LD, Ctrl,2}}(t) \end{pmatrix}, \begin{pmatrix} n_{\text{LD, Ctrl,3}}(t) \\ r_{\text{LD, Ctrl,3}}(t) \end{pmatrix}, \begin{pmatrix} n_{\text{MD, RSL3,1}}(t) \\ r_{\text{MD, RSL3,1}}(t) \end{pmatrix}, \dots \right\}.$$

We performed parameter inference using all replicates $D_{\text{rep}}$ obtained from 12-well experiments. The dataset comprises three independent replicates (labelled $j$ before) initiated at low (LD), medium (MD), and high (HD) cell density under two conditions (Ctrl and RSL3). The dynamics were tracked over 13 time points over a 24-hour period with intervals of 2 hours. The fit was performed with a single set of parameters for all replicates and seeding conditions, with only the parameter $r_0$ allowed to vary between the Ctrl and RSL3 conditions.

##### 3.4 Fixed parameters and variable initial conditions

For the optimisation, we have fixed $K = 100\%$ confluence as the maximum carrying capacity. Similarly, we set the Hill coefficient $h = 2$ from here onwards as it is a minimum value at which non-linear effects can cause bistability for $f = 0$, see also Appendix 4). For the ROS feedback, the Hill coefficient $l = 3$ is chosen based on Co et. al[1] where a positive feedback loop of ROS biochemistry is modelled as a hyperbolic function with a Hill coefficient of 3.

In order to solve the differential equation numerically, initial values $n(t = 0)$ and $r(t = 0)$ are set, which determine the course of the populations deterministically. Rather than fixing them a priori, we allow the optimisation of these initial conditions of a trajectory. We thus account for noise in the data at $t = 0$, since the population may need time to establish itself in the vessel. On this basis, we can find the most suitable parameter set $p$ by minimizing the loss function $L(p)$, as explained in section 3.2.

Importantly, we introduce a penalty to our loss function to favour solutions in which trajectories that start at high densities (HD) grow reliably to confluence. Using a tolerance of $n(t = 0) = 60\%$, we define a threshold density for which this steady state condition must apply. This cost is both technically and biologically motivated, as the equations also allow solutions in which high cell densities reach almost confluence but will then go extinct after a long period of time. However, such long-term dynamics are not observed within our experimental observation window and do not adequately reflect our data. Furthermore, we add a cost for initial conditions $n(t = 0) > 100\%$ during the fitting process. These are mathematically possible, but not suitable for experimental data that is spatially limited.

##### 3.5 Fit results

We performed a grid search over the parameter space by using multiple combinations of initial values in the parameter space for the optimisation to identify the set that minimises the loss function. Initial conditions for each time series, $(n_{\text{pred},j}(t = 0), r_{\text{pred},j}(t = 0))$, were always initialised directly from the corresponding experimental measurements, $(n_{\text{data},j}(t = 0), r_{\text{data},j}(t = 0))$. For the 12-well time series dataset $D_{\text{rep}}$ the parameters were estimated as follows: the time dependent rates were $a = 1.6 \cdot 10^{-2}\text{hour}^{-1}$, $b = 6.6 \cdot 10^{-2}\text{hour}^{-1}$ and $\beta = 3.1 \cdot 10^{-3}\text{hour}^{-1}$, while $\alpha = 9.9 \cdot 10^{-5}$, $r_1 = 1.4 \cdot 10^{-3}$ and the condition-specific values $r_{0,\mathrm{Ctrl}} = 1.8 \cdot 10^{-3}$ and $r_{0,\mathrm{RSL3}} = 5.5 \cdot 10^{-4}$ are expressed in units of the green-to-orange fluorescence ratio. Similarly, the production rate $f = 2.3 \cdot 10^{-4}\mathrm{hour}^{-1}$ is given units of the green-to-orange fluorescence ratio per hour. The lipid ROS-related parameters and measurements could, in principle, be calibrated to absolute numbers of lipid ROS molecules per cell. The initial conditions of the individual trajectories, which are optimised over the course of the fitting process, are not listed. The corresponding plots are shown in Figures 5 and 6. To identify the underlying basins of attraction, we simulated trajectories from a grid of initial conditions $(n(t = 0), r(t = 0))$ using the fitting parameters. The ROS initial conditions were sampled log-uniformly. For each initial condition, we numerically integrated the differential equations and checked whether the trajectory converged within a certain tolerance towards confluence with minimised lipid ROS values $(K, 0)$ or not. The resulting binary map shows the ferroptosis-insensitive basin of attraction (blue) and the ferroptosis-sensitive basin of attraction (orange) in the $n$-$r$ phase space and the phase space boundary.

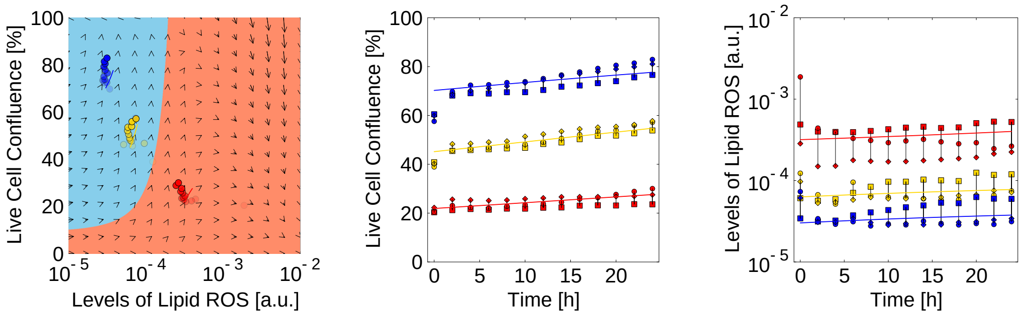

Figure 5: **Replicates of the time series data compared to the least-squares estimate (Ctrl condition).** The data from several replicates are shown as scatter points and are linked with a line. The fits are shown as thick lines. For a clearer illustration, a single representative fitting curve was drawn per seeding density, with the initial condition being the median of the fitted initial conditions. Red, yellow, and blue are chosen for LD, MD, and HD, respectively.

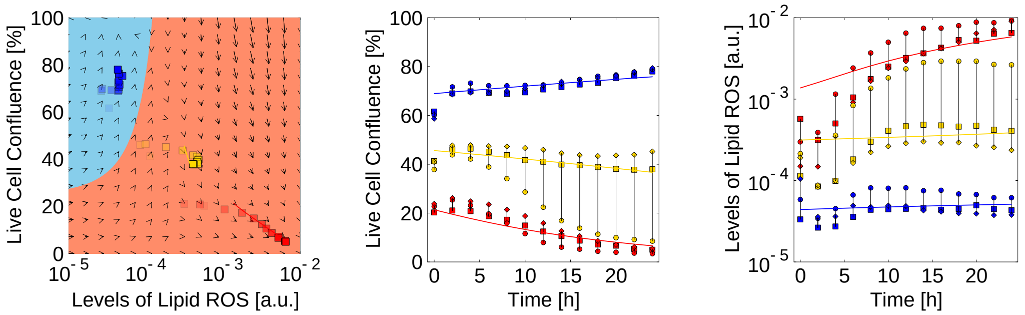

Figure 6: **Replicates of the time series data compared to the least-squares estimate (RSL3 condition).** The data from several replicates are shown as scatter points and are linked with a line. The fits are shown as thick lines. For a clearer illustration, a single representative fitting curve was drawn per seeding density, with the initial condition being the median of the fitted initial conditions. Red, yellow, and blue are chosen for LD, MD, and HD, respectively.

#### 4 Addendum: Fixed points without positive feedback

Fixed points are found by solving the system of equations

$$\dot{n}(t) = 0, \tag{4}$$
$$\dot{r}(t) = 0 \tag{5}$$

for $f = 0$. We calculate two fixed points for $h = 2$ and find two stable fixed points that allow a bistability analysis and possible interpretation of our experimental observations

$$FP = (n^*, r^*) = (K, 0)\,, \tag{6}$$
$$FP = (n^*, r^*) = \left(K\left(1 - \frac{b}{2a}\left(1 - \frac{1}{\gamma}\sqrt{\gamma^2 - 1}\right)\right), r_0\left(\gamma - \sqrt{\gamma^2 - 1}\right)\right), \tag{7}$$
$$FP = (n^*, r^*) = \left(K\left(1 - \frac{b}{2a}\left(1 + \frac{1}{\gamma}\sqrt{\gamma^2 - 1}\right)\right), r_0\left(\gamma + \sqrt{\gamma^2 - 1}\right)\right), \tag{8}$$
$$FP = (n^*, r^*) = \left(0, \frac{a\alpha}{\beta}\right). \tag{9}$$

with $\gamma \equiv \frac{\alpha b}{2\beta r_0}$. Here, the first solution represents a global stable fixed point. This point represents a population that, in the long-time limit, succeeds to grow to confluence and minimises the lipid ROS level per cell. The last fixed point represents a semi-stable solution, that only is approached for $n(t = 0) = 0$ or under specific parameter combinations. Such parameter sets are if the third fixed point converges towards the fourth. Then, for a bistable system, the population either goes extinct $n = 0$ or is driven towards the carrying capacity $n = K$. In between the two stable fixed points we find a saddle point, the second solution, dividing the phase space $n, r$ in two basins of attraction. The fixed points can disappear with a saddle-node bifurcation, e.g. with a critical parameter, this could be $\beta$, the bistable system could collapse into a monostable system where only $n = K$ remains as a solution.

### References

[1] Co, H. K. C.; Wu, C.-C.; Lee, Y.-C.; Chen, S.-h. *Nature* **2024**, *631*, 654–662.
